# Pathogen-specific structural features of two key players in *Candida albicans* morphogenetic switch

**DOI:** 10.1101/2022.08.23.504951

**Authors:** José A Manso, Arturo Carabias, Zsuzsa Sárkány, José M de Pereda, Pedro José Barbosa Pereira, Sandra Macedo-Ribeiro

## Abstract

Ras-like protein 1 (CaRas1) is a key regulator of the switch between the yeast and hyphal forms of *Candida albicans*, a feature associated with pathogenesis. CaRas1 is activated by the guanine nucleotide exchange factor (GEF) CaCdc25, triggering hyphal growth-related signaling pathways through its highly conserved GTP-binding domain (G-domain). An important function in hyphal growth has also been proposed for the long hypervariable region downstream of the G-domain of CaRas1, whose unusual content of polyQ stretches and Q/N repeats make CaRas1 unique within Ras-family proteins. Despite its biological importance, both the structure of CaRas1 and the molecular basis of its activation by CaCdc25 remain unexplored. Here, we show that CaRas1 displays an elongated shape and that its hypervariable region contains helical structural elements with intramolecular coiled-coil propensity and limited conformational flexibility. Functional assays revealed that CaRas1 activation by CaCdc25 is highly efficient, with 5-to 2000-fold higher activity levels than reported for human GEFs. In addition, the threedimensional structure of the catalytic region of CaCdc25, together with the structural characterization of CaRas1/CaCdc25 complexes, unveiled a specific region located in the α-helical hairpin of CaCdc25, critical for CaRas1 activation, where negatively charged substitutions reduce its activity. The unique structural features of the low complexity region of CaRas1 and the distinctive properties of CaRas1 activation by CaCdc25, common in the homologous proteins from CTG-clade species, uncover novel strategies to target key virulence factors in human-infecting fungal pathogens.

## Introduction

*Candida albicans*, part of the commensal microbiota of most individuals, is an important opportunistic human pathogen and can cause severe infections in immunocompromised patients (Beck-Sagué and Jarvis, 1993). *C. albicans* belongs to the CTG-clade of *ascomycetes*, which includes a high number of opportunistic pathogenic species that use an alternative genetic code and translate the CUG codon as a serine instead of leucine (Rossignol et al., 2008; Turner and Butler, 2014). The remarkable adaptability of *C. albicans* to highly diverse host niches (Nikou et al., 2019) is underscored by its phenotypic plasticity, i.e., the discrete phenotypes adopted in response to varying environmental cues (Scaduto and Bennett, 2015). The commensal-to-pathogen transition of *C. albicans* is associated with its ability to interconvert between yeast and hyphal morphologies (Lo et al., 1997; Mitchell, 1998; Peters et al., 2014). The activation of hyphae-specific genes is directly mediated by the cyclic adenosine monophosphate (cAMP) and the mitogen-activated protein kinase (MAPK) signaling pathways, which are regulated by the Ras-like protein 1 (CaRas1) and the cell division control protein 25 (CaCdc25) (Leberer et al., 2001; Pentland et al., 2017).

CaRas1 is composed of a highly conserved GTPase domain (G-domain) followed by a hypervariable region. The long hypervariable region of CaRas1 represents a major difference between yeast (~120 residues) and human (~20 residues) Ras proteins. Further, the presence of stretches of low complexity segments in the hypervariable region of CaRas1, including polyglutamine (PolyQ) repeats and a Q/N rich stretch, also present in other members of the CTG-clade, makes it unique among Ras-family proteins (Figure S1). The hypervariable region of CaRas1 also contains a C-terminal CCaaX membrane association motif, whose two cysteine residues have been found to be farnesylated and palmitoylated (Piispanen et al., 2011), in agreement with the localization of CaRas1 to membranes, as observed for many Ras GTPases. Interestingly, cleavage of the last 67 amino acid residues of CaRas1 at its hypervariable region has been proposed as a mechanism modulating CaRas1 signaling and, consequently, hyphal growth (Piispanen et al., 2013).

In yeast cells, CaRas1 is generally in an inactivated, GDP-bound form (CaRas1-GDP). CaCdc25 activates CaRas1, converting it to CaRas1-GTP and switching on the signaling pathway associated with the hyphae-specific genes (Lai et al., 1993; Maidan et al., 2005; Pentland et al., 2017). The activation of CaRas1 by CaCdc25 is the primary mechanism by which D-glucose induces morphogenesis (Maidan et al., 2005; Pentland et al., 2017; Rolland et al., 2000). In fact, deletion of either CaRas1 or CaCdc25 results in hyphal defects (Parrino et al., 2017). CaCdc25 (~150 kDa) is a Ras-guanine nucleotide exchange factor (GEF) that contains a C-terminal catalytic region, consisting of the tandem Ras-exchange motif domain-catalytic domain (REM-CAT), which catalyzes the exchange of CaRas1-bound GDP to GTP.

Despite its biological relevance, no experimental structural information is available for the full-length CaRas1 and the arrangement of the G-domain and the unique low complexity hypervariable region, which is predicted to be partially disordered. Also, the elucidation of the molecular basis for the activation of CaRas1 by CaCdc25 would provide crucial information for the design of new drugs to fight this fungal human pathogen.

In order to elucidate the structural features of this unique Ras-family protein, we determined the solution structure of full-length CaRas1 and of truncated variants at low resolution, using a combination of small-angle X-ray scattering (SAXS), homology modelling, secondary structure prediction and circular dichroism (CD). Our results disclosed a hitherto unknown elongated conformation for CaRas1, with an unexpected limited flexibility of its singular hypervariable region. In addition, the crystal structure of the catalytic region of CaCdc25 (REM-CAT) and the solution structures of the CaRas1/REM-CAT complex provided a valuable insight into the molecular details of the CaRas1/REM-CAT interaction, unveiling a *Candida-specific* region of CaCdc25 required for the activation of CaRas1, strikingly conserved in most common fungal pathogens and largely divergent in mammalian GEF proteins.

## Results

### The domain organization of CaRas1 is unique within the family of Ras-like proteins

CaRas1 is composed by a highly conserved G-domain, followed by an exclusive hypervariable region that contains low complexity sequence segments, including PolyQ repeats and a Q/N rich stretch (Figure 1*A*), a unique feature conserved in Ras-family proteins from various of the CTG-clade opportunistic fungal pathogens. The hypervariable region of CaRas1 is annotated as disordered in the MobiDB database (Piovesan et al., 2021), which incorporates consensus analysis from several disorder prediction servers. Interestingly, three independent secondary structure predictors - PsiPred, Porter and JPred4 - assign helical elements within the polyQ and Q/N rich stretches of the hypervariable region (Figure 1*A*), in line with AlphaFold2 predictions (Varadi et al., 2022). This is not completely unexpected since a role has been described for polyQ regions in the formation of coiled-coil structures, particularly if preceded by a region with high helical propensity (Fiumara et al., 2010; Mier and Andrade-Navarro, 2020).

**Figure 1.**
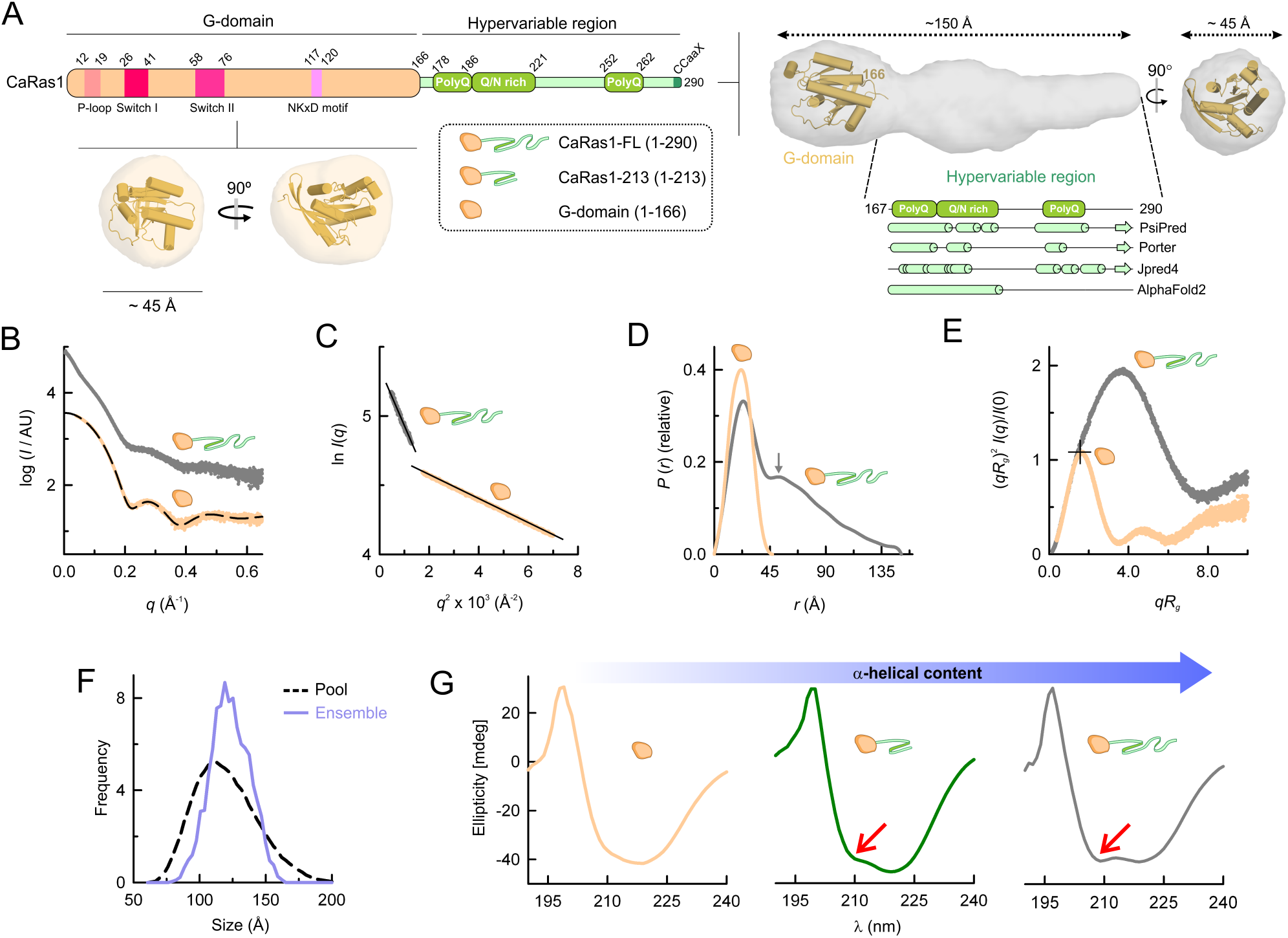
The hypervariable region of CaRas1 displays an unexpected architecture. A) Schematic domain organization of CaRas1, comprising the G-domain (residues 1-166) and the hypervariable region (residues 167-290). Important structural elements of the G-domain - nucleotide exchange P-loop, Switches I and II and NKxD motif - are indicated and the polyQ and Q/N-rich segments are highlighted within the hypervariable region. Schemes of the different constructs used in this work are provided in the dotted box. The SAXS-derived molecular envelopes for the CaRas1 G-domain and for the full-length protein are shown as semi-transparent surfaces, below and to the right of the domain organization scheme, respectively. A cartoon representation (orange) of the G-domain model is docked into the envelopes. Below the envelope of full-length CaRas1, predicted secondary structural elements for the hypervariable region are represented, suggesting that the polyQ and Q/N-rich regions are mainly helical. B) Experimental SAXS profiles extrapolated to infinite dilution of CaRas1-FL (dark-grey) and G-domain (orange). Curves are offset on the log scale for readability; the same data color scheme is used in C–G. The scattering calculated for the atomic model of the G-domain shown in panel A is represented as a black dashed line. C) Guinier plots of the scattering data shown in B. D) *P*(*r*) functions determined from the scattering data shown in B. The maximum for a second peak in CaRas1-FL is indicated by a grey arrow. E) Dimensionless Kratky plots. The crosshair indicates the expected position of the theoretical maxima of the plot for spherical compact particles (*qR*_g_ ~ 1.732 and (*qR*_g_)^2^*I*(*q*)/*I*(0) ~ 1.104). F) Ensemble optimization methods (EOM) analysis of the flexibility of the hypervariable region. Frequency of size distributions in a pool of 10,000 models (black dashed line) with random orientations of the hypervariable region, and in the selected ensemble that fits the SAXS data of CaRas1-FL (blue line). G) Circular dichroism spectra of CaRas1 G-domain, CaRas1-213 (green) and CaRas1-FL. The negative band at 209 nm (red arrows) observed when the hypervariable region is present in CaRas1, together with that at 220 nm are spectral signatures for α-helical content.

The structural properties of the hypervariable region within CaRas1 and its relative position with respect to the globular G-domain were evaluated using SAXS (Figure 1*A-F*, Table S1) and two CaRas1 constructs, the full-length protein (CaRas1-FL, 1-290) and the isolated G-domain (1-166). Both CaRas1 constructs were monodisperse and monomeric in solution (Figure 1*C*, Table S1). The distance distribution function (*P*(*r*)) for the G-domain has a bell shape characteristic of globular particles with a peak at ~23 Å and a maximum dimension (*D*_max_) of ~45 Å (Figure 1*D*) in good agreement with a homology model for the G-domain (Figure S2, Figure 1*A,B*) that could be easily docked in a low-resolution envelope obtained by *ab initio* methods from the experimental SAXS data (Figure 1*A*). Interestingly, the high resolution of the SAXS data allowed to distinguish between similar homology models and suggests that the highly dynamic switch II region of G-domain displays an α-helical structure in CaRas1 (Figure S2). When the hypervariable region is present, a second peak appears at ~52 Å in the *P*(*r*) function (Figure 1*D*), suggesting a certain degree of rigidity and the presence of structured elements in that region of CaRas1-FL. Interestingly, the low-resolution envelope for CaRas1-FL, which is an average of 15 independent *ab initio* models (Table S1), displays an elongated shape (Figure 1*A*). The homology model for the G-domain could be docked in one end of this envelope, allowing the assignment of an adjacent volume to the hypervariable region (Figure 1*A*). The differences between CaRas1-FL and its isolated G-domain are further highlighted in the dimensionless Kratky plot of the scattering data (Receveur-Bréchot and Durand, 2012), which displays a bell-shaped peak with a maximum at 1.1 for the G-domain, indicating that it is a compact and spherical particle (Figure 1*E*). When the hypervariable region is present, a bell-shaped and sharp peak is also observed in this plot, yet the position and amplitude of its maximum deviates largely from the value expected for spherical particles, in line with the highly anisometric shape of CaRas1-FL (Ortega et al., 2016) (Figure 1*E*). The overall shapes of the bimodal *P*(*r*) function and the dimensionless Kratky plot for CaRas1-FL, are characteristic of a protein composed by relatively rigid two-bodies (Sethi and Ylänne, 2014), suggesting that the hypervariable region is not entirely disordered.

In order to better understand the structure and flexibility of the hypervariable region, an ensemble fitting method was employed (Bernadó et al., 2007). A pool of 10,000 CaRas1-FL structures was generated, sampling exhaustively the potential orientations of the hypervariable region, assuming it to be fully flexible. The minimal ensemble that better approximated the experimental SAXS data was mainly populated by conformations with a narrower size distribution than the entire size range of the pool (Figure 1*F*), further underscoring that the hypervariable region is not fully flexible. Besides, from this analysis a Rsigma value of 0.61 was calculated and values < 1.0 for this parameter are indicative of limited flexibility in the system. In agreement, when the secondary structure of three CaRas1 truncation constructs with variable lengths of the hypervariable region (Figure 1*A*) was analyzed by CD, the unanticipated ordered structure of this C-terminal tail came to light. The CD spectra show an increase in the α-helical content when the hypervariable region is present (Figure 1*G*). Importantly, the CD spectrum of CaRas1-FL displays minima at 220 nm and 209 nm, with a 220 nm/209 nm ratio of 1.00, supporting the presence of a coiled-coil structure in the hypervariable region. In the presence of 50% (v/v) trifluoroethanol (TFE), a strong a-helix stabilizer that disrupts coiled-coil formation, there is a shift of the 220 nm/209 nm ratio to 0.88 (Figure S3), which is more suggestive of isolated helices (Zhou et al., 1992).

### Structure of the catalytic region of the CaRas1 guanine exchange factor CaCdc25

The crystal structure of the catalytic region of the CaRas1 guanine nucleotide exchange factor CaCdc25, which catalyzes the exchange of the bound nucleotide and therefore the conversion of CaRas1-GDP to CaRas1-GTP, was determined at 2.45 Å-resolution by X-ray crystallography (Figure 2*A-F*, Table S2). The protein crystallized in the orthorhombic space group *P*2_1_2_1_2_1_ with two molecules of CaCdc25 in the asymmetric unit (AU) that are very similar (Figure S4*A*,*B*). The monomers superpose with a root mean square deviation (rmsd) of 1.0 Å for 384 *C*α atoms. A structural homology search with DALI (Holm, 2020) showed that despite low amino acid sequence conservation, the structure of catalytic region of CaCdc25 is quite similar to that of human GEFs containing Cdc25 homology domains and that act on the Ras family (four matches with Z score > 20; Figure S4*C*), and consists of two well-differentiated domains: the REM domain (residues 883-1035, β-strands β1 and β2 and a-helices α1-α10) and the CAT domain (residues 1039-1306, β-strands βA and α-helices αA-αQ) (Figure 2*A*). In particular, the individual structures for the CAT domain in the five molecules are more conserved than for the REM domain (Figure S4*D*). After pairwise superimposition of the five structures of the CAT domain, the rmsd for 286 *C*α atoms ranged from 3.7 to 4.7 Å. In contrast, the five copies of REM were superposed with a rmsd for 179 *C*α atoms between 6.3 and 7.9 Å.

**Figure 2.**
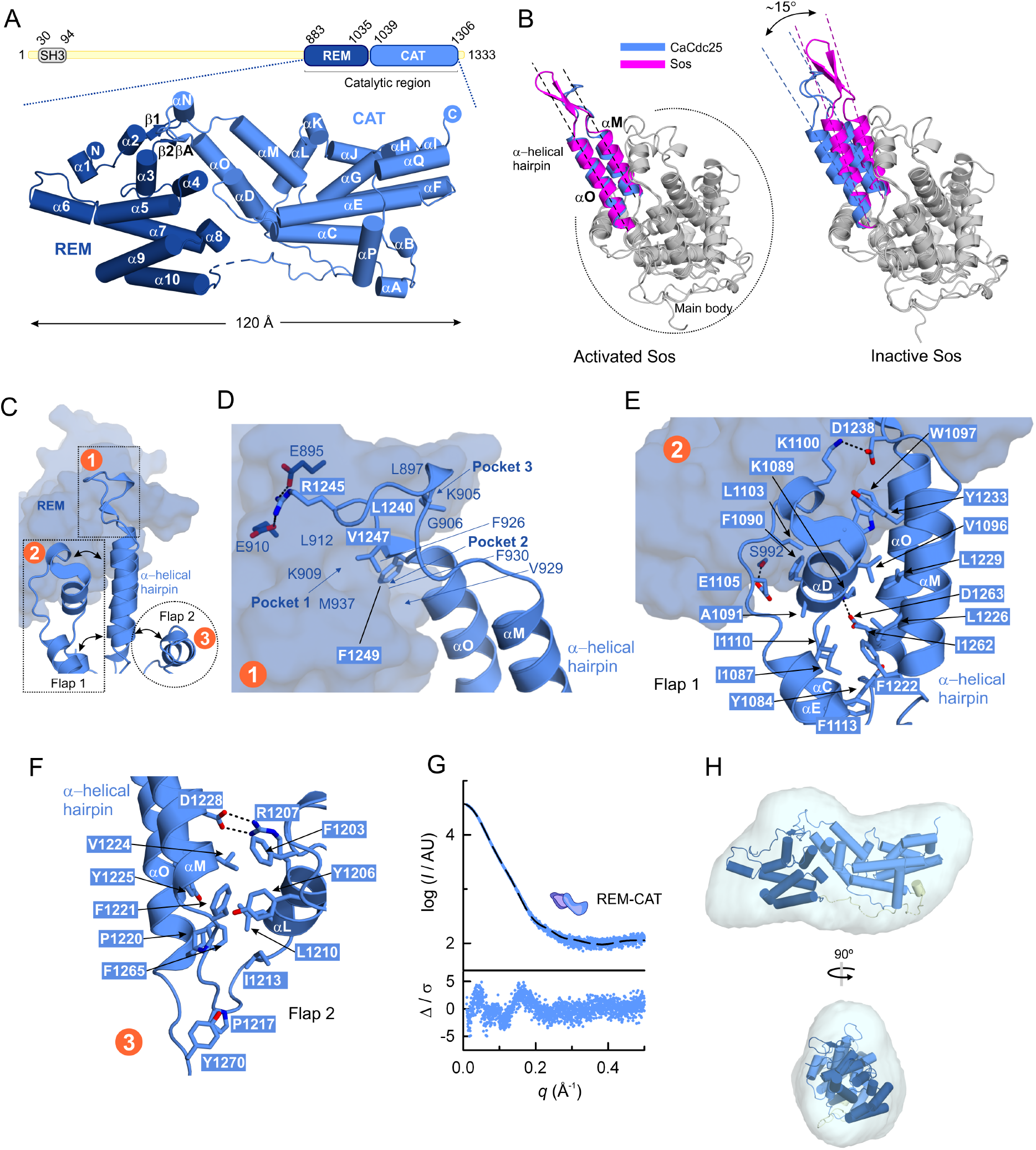
Structure of the catalytic region of CaRas1-guanine nucleotide exchange factor CaCdc25. A) Domain organization diagram (top) and cartoon representation (bottom) of the crystal structure of the catalytic region of CaCdc25 at 2.45 Å resolution. The REM (dark blue) and CAT (light blue) domains are labelled and the N- and C-termini are indicated in filled circles. B) Structural superposition of the main body of the CAT domain of CaCdc25 with the active (PDB entry 1NVW (Margarit et al., 2003)) and inactive (PDB entry 2II0 (Freedman et al., 2006b)) forms of Sos showing that the α-helical hairpin in the CaCdc25 structure is poised for activity. C) Overview of the three main interacting regions (numbered in orange circles) that maintain the α-helical hairpin in an activating position, detailed in panels D-F. D) Interaction of the loop connecting the two main helices of the α-helical hairpin (main interacting residues shown as sticks) with the REM domain (transparent surface). E) Interaction of the α-helical hairpin with the flap1 region (main interacting residues shown as sticks) and with the REM domain (transparent surface). F) Interaction of the α-helical hairpin with the flap2 region (main interacting residues shown as sticks). G) Experimental SAXS profile of the catalytic region of CaCdc25, extrapolated to infinite dilution (upper plot). The scattering calculated for the atomic structure of the tandem REM-CAT of CaCdc25, shown in A, is represented as a black dashed line. Error-weighted residual difference plot Δ/σ = [*I*_exp_(*q*) - *cI*_mod_(*q*)]/ σ (q) versus *q* (lower plot). H) Crystal structure of CaCdc25 (cartoon representation) docked into the SAXS-derived molecular envelope (translucent surface), which is the average of 15 independent bead models. Two orthogonal views are shown.

The α-helical hairpin is a key element of the CAT domains for the Ras nucleotide exchange in GEFs (Gasper and Wittinghofer, 2020). In CaCdc25 the α-helical hairpin is formed by the two antiparallel helices α**M** and α**O** (residues 1222-1265) (Figure 2*B*), which protrude from the main body of the CAT domain. The relative orientation of the α-helical-hairpin is critical for the activity of GEFs (Freedman et al., 2006a). In CaCdc25, the hairpin assumes an orientation similar to that observed in the structure of the active state of the human HRas GEF Sos (Figure 2*B*), suggesting that the CAT domain of CaCdc25 is also in the active conformation in the crystal structure. The active state-like orientation of the α-helical hairpin of CaCdc25 is stabilized mostly by hydrophobic interactions with three main regions (Figure 2*C*): **1)** the REM domain in which α-helical hairpin residues L1240, V1247 and F1249 occupy three hydrophobic grooves formed by K909, L912 and M937 (pocket 1), F926, V929 and F930 (pocket 2), and L897, K905 and G906 (pocket 3) (Figure 2*D*), **2)** a region called flap 1 (residues 1080-1115) that encompasses helix α**D** and part of helices α**C** and α**E**, and that stabilizes helix α**M** of the hairpin (Figure 2*E*) and **3)** the region called flap 2 (residues 1195-1215), which includes part of helix α**L** (Figure 2*F*). In addition to these hydrophobic interactions, typical of the GEF family, the α-helical hairpin of CaCdc25 is further stabilized by three unique polar contacts, which are not observed in the structures of GEFs listed in Figure S4*C*: R1245 makes ionic interactions with E895 and E910 from the REM domain (Figure 2*D*); D1238 interacts with K1100 from the flap 1 region; and E1105 from flap 1 interacts with S992 from the REM domain (Figure 2*E*). Importantly, comparison with human GEFs revealed only partial conservation of the residues mediating the activation of Ras (Figure S5). The two important residues located in the α-helical hairpin that impede nucleotide binding in the human GEF Sos1, L938 and E942, are replaced by T1230 and H1234, respectively, in CaCdc25, which, together with the observed differences in the distribution of surface electrostatic potential in the G-domain interacting region (Figure S6), suggests the REM-CAT tandem of CaCdc25 as a potential target for specifically inhibiting Ras1-activation in *Candida albicans* without affecting other human GTPases.

Finally, the solution structure of catalytic region of CaCdc25 is in excellent agreement with the crystallographic model (Figure 2*G*, Table S1), which unsurprisingly docks well into the *ab initio* molecular envelope derived from the SAXS data (Figure 2*H*).

### Solution structures of the CaRas1/REM-CAT complexes

When mixed in equimolar amounts, the catalytic region of CaCdc25 and either CaRas1-FL or its isolated G-domain form complexes detectable by size-exclusion chromatography (SEC) (Figure S7). SAXS analysis (Figure 3*A*) of these complexes revealed monodispersity of the samples (Figure 3*B*) and 1:1 stoichiometry, as deduced from the estimated masses (Table S1). A *P*(*r*)-derived *D*max of ~120 Å for the G-domain/REM-CAT complex (Figure 3*C*) is in good agreement with an atomic model for the complex (Figure 3*D*), which was generated after superposition of our REM-CAT and G-domain structures onto the structure of Sos1 complexed with human HRas (PDB ID 1BKD (Boriack-Sjodin et al., 1998)). The theoretical scattering calculated for the model of G-domain of CaRas1 in complex with the catalytic region of CaCdc25 reproduces well the experimental data (Figure 3*A*). Further, this model also fitted nicely the low-resolution molecular envelope (Figure 3*D*), suggesting that the interaction between these proteins globally resembles that observed for human HRas and Sos1.

**Figure 3.**
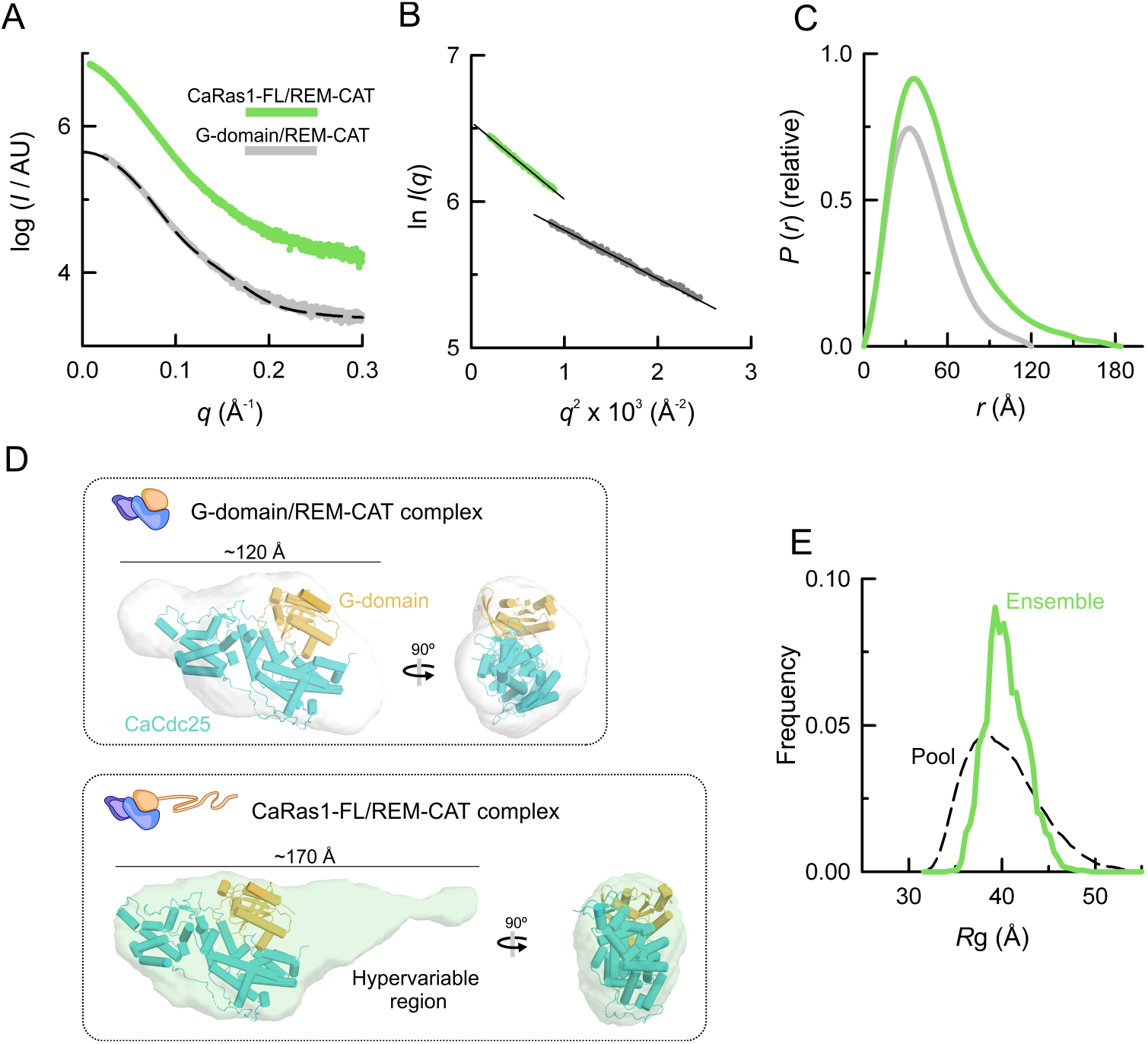
Structures of the CaRas1/REM-CAT complexes in solution. A) Experimental SAXS profiles extrapolated to infinite dilution of the complexes of the catalytic region of CaCdc25 with CaRas1-FL and CaRas1 G-domain. Curves are offset on the log scale to improve readability. The scattering curve calculated for the atomic model of the complex shown in D (upper panel) is plotted as a black dashed line. B) Guinier plots of the scattering data shown in A. C) *P*(*r*) functions determined from the scattering data shown in A. D) Cartoon representation of the REM-CAT/CaRas1 G-domain complex docked into the SAXS-derived molecular envelopes for the complexes of CaCdc25 with the CaRas1 G-domain (upper panel) and CaRas1-FL (lower panel), shown as grey and pale green semitransparent surfaces, respectively. The low-resolution structures are the average of 15 independent bead models. Two orthogonal views are shown. E) Ensemble optimization methods (EOM) analysis of the flexibility of the hypervariable region in the CaRas1-FL/REM-CAT complex. Size frequency distributions in a pool of 10,000 models (black dashed line) with random orientations for the hypervariable region, and in the selected ensemble that fits the SAXS data of CaRas1-FL (green line). As for the isolated CaRas1-FL, a Rsigma value < 1.0 (0.55) is obtained and is indicative of a system with limited flexibility.

The *ab initio* reconstruction of the envelope for the complex with CaRas1-FL reveals a more extended shape than for the G-domain (Figure 3*C,D*). The CaRas1 G-domain/REM-CAT complex model could be docked in this envelope, revealing the relative position of the hypervariable region of CaRas1-FL (Figure 3*D*) that appears to be in an elongated conformation, as was observed for the isolated CaRas1. Further, the hypervariable region displays limited flexibility in the context of the complex with the catalytic region of CaCdc25 (Figure 3*E*), suggesting the absence of a conformational change in this region upon complexation. This experimental low-resolution envelope nicely resembles the predicted complex models by AlphaFold2 (Figure S8), in which the hypervariable region of CaRas1, part predicted as a single and long helix (see “Discussion” section), seems to be not involved in the interaction, suggesting that it is unlikely to interfere in the CaCdc25-modulated nucleotide exchange reaction.

ITC data confirmed that the complex formation was unaffected by the presence of the hypervariable region of CaRas1 and revealed that the catalytic region of CaCdc25 interacted with CaRas1-FL or CaRas1 G-domain with μM affinities (*K*_d_ = 9 μM) (Figure 4*A*). As suggested by the SAXS data, CaCdc25 forms complexes with CaRas1 with a 1:1 stoichiometry (Figure 4*A*). Also, varying the length of the hypervariable region did not affect the nucleotide exchange activity of the catalytic region of CaCdc25 (Figure 4*B*), with similar mean rate constants (*k*_obs_) observed for three different CaRas1 constructs (Figure 4*C*), further suggesting that the hypervariable region has no effect on the nucleotide exchange activity. Interestingly, the activity observed for the catalytic region of CaCdc25 was considerably higher (from 5-to 2000-fold) (Table S3) than those previously reported for other GEFs, under similar experimental conditions (Carabias et al., 2020; Popovic et al., 2013, 2016) underscoring the efficiency of CaRas1 activation by this GEF.

**Figure 4.**
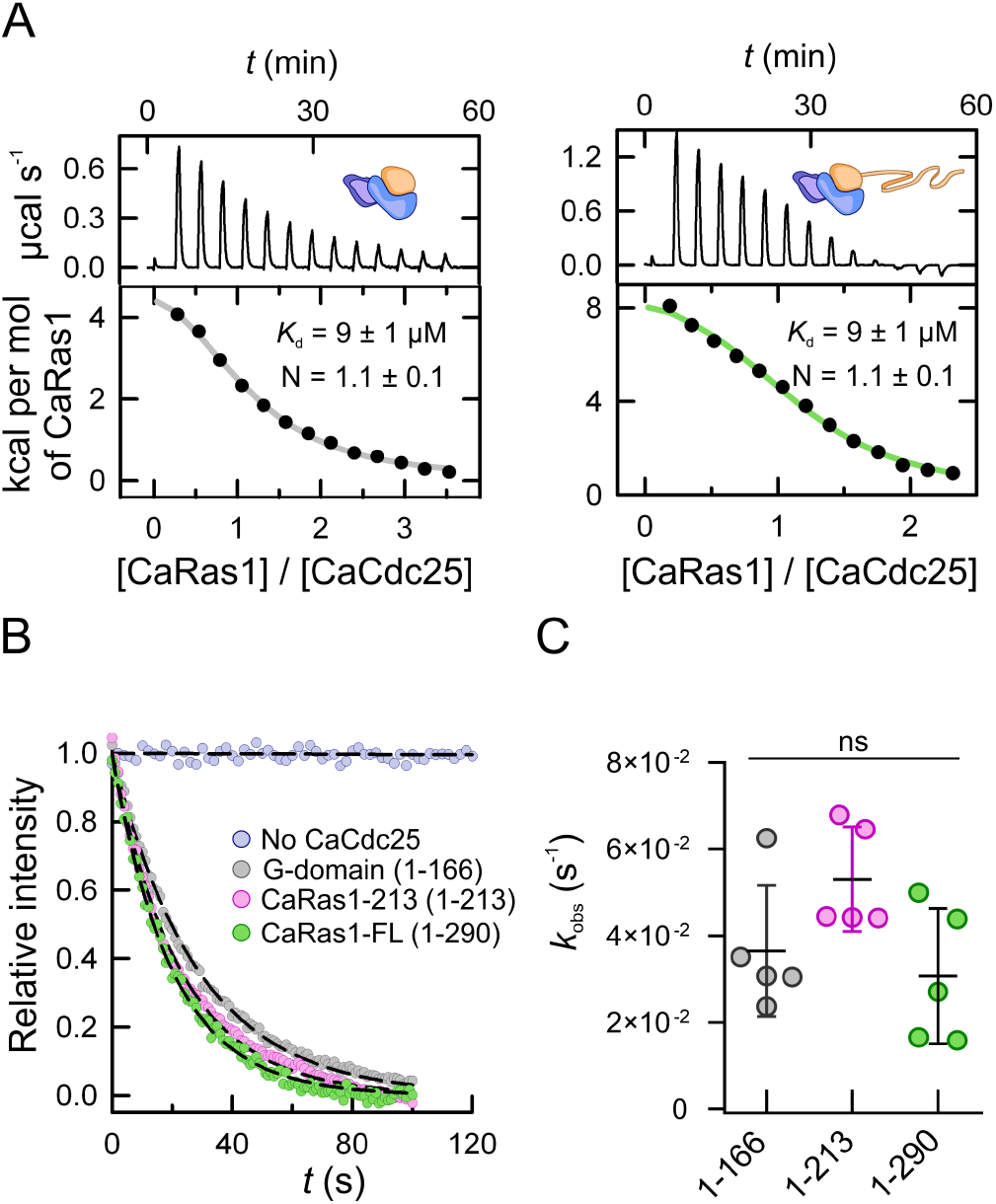
Characterization of the interaction of the catalytic region of CaCdc25 with CaRas1 and evaluation of the effect of the hypervariable region on the activity. A) Thermograms (upper panels) and binding isotherms (lower panels) of the binding of the REM-CAT tandem of CaCdc25 to the G-domain of CaRas1 (left) and to CaRas1-FL (right). The data were analyzed fitting a model assuming one binding site in the catalytic region of CaCdc25 for CaRas1 G-domain (grey line) and CaRas1-FL (green line). The presence of the hypervariable region does not affect the affinity, suggesting that it does not impact the recognition of CaCdc25. B) Representative nucleotide exchange reactions. Mant-dGDP loaded CaRas1 G-domain, CaRas1-213 and CaRas1-FL were used at 200 n*M* in absence and in presence of 100 n*M* CaCdc25. C) The *k*_obs_ values for the nucleotide exchange reactions were obtained by fitting a single exponential decay model (dashed lines in H) to the experimental data. All kinetics were performed in quintuplicate. Lines represent mean ± SD of the replicates. The differences observed for the different constructs were not significant (ns).

### Negatively charged substitutions at the uniquely conserved H1234 in the α-helical hairpin decrease the activity of CaCdc25

The GDP/GTP exchange reaction in Ras proteins is stimulated through their interaction with the CAT domain of GEFs, a mechanism well conserved from yeast to humans (Chardin et al., 1993; Créchet et al., 1990). Once the GEF binds the GDP-bound, inactive Ras, the α-helical hairpin of the CAT domain has a crucial role in nucleotide release and consequently in Ras activation. In particular, the side chains of residues from this hairpin modify the chemical environment of the nucleotide binding site, promoting GDP release (Boriack-Sjodin et al., 1998). Subsequently, GTP binding to Ras not only activates it but also disrupts the Ras-GEF complex. In the Sos1-HRas complex L938 and E942 from Sos1 (the latter also present in the human GEFs RASGRF1, EPAC2 and RASGRP1), are key residues for nucleotide release, affecting the binding of Mg^2+^ and of the α-phosphate group of the nucleotide, respectively. In order to further validate our CaRas1/REM-CAT complex model (Figure 3D), we evaluated the effect of single residue substitutions at position 1234 of the α-helical hairpin, which is structurally equivalent to the important E942 in Sos1 (Figure 5A), in the CaRas1 nucleotide exchange activity. Interestingly, there was a four-fold reduction in the unusually high activity of the catalytic region of CaCdc25 (*k*_obs_ = (4 ± 2) 10^-2^ s^-1^) when H1234 was replaced by a negatively charged residue (*k*_obs_ = (1.2 ± 0.1) 10^-2^ s^-1^ for H1234E and *k_obs_ =* (0.90 ± 0.05) 10^-2^ s^-1^ for H1234D), while replacement with an uncharged amino acid did not significantly alter the activity (*k*_obs_ = (3.3 ± 0.4) 10^-2^ s^-1^ for H1234A) (Figure 5*B,C*). Of note, sequence conservation analysis with ConSurf (Ashkenazy et al., 2016) led to the identification of 150 homologues of CaCdc25 in fungi. Within this group of homologous proteins, a histidine residue is strictly conserved in a position equivalent to that of H1234 in CaCdc25 only in ten annotations, all of them belonging to the CTG-clade that contains most of the pathogenic *Candida* species (Gabaldón et al., 2016; Turner and Butler, 2014), while alanine and valine are found in the *Saccharomyces cerevisae* and *Candida glabrata* homologues, respectively (Figure S9). The remaining sequences have mostly a negatively charged residue (129 have an aspartate and 3 a glutamate) in the position equivalent to H1234 of the catalytic region of CaCdc25, and belong to non-pathogenic fungi. In summary, H1234 in the α-helical hairpin is a key residue for the GEF activity of CaCdc25 that is unique to CTG-clade species.

**Figure 5.**
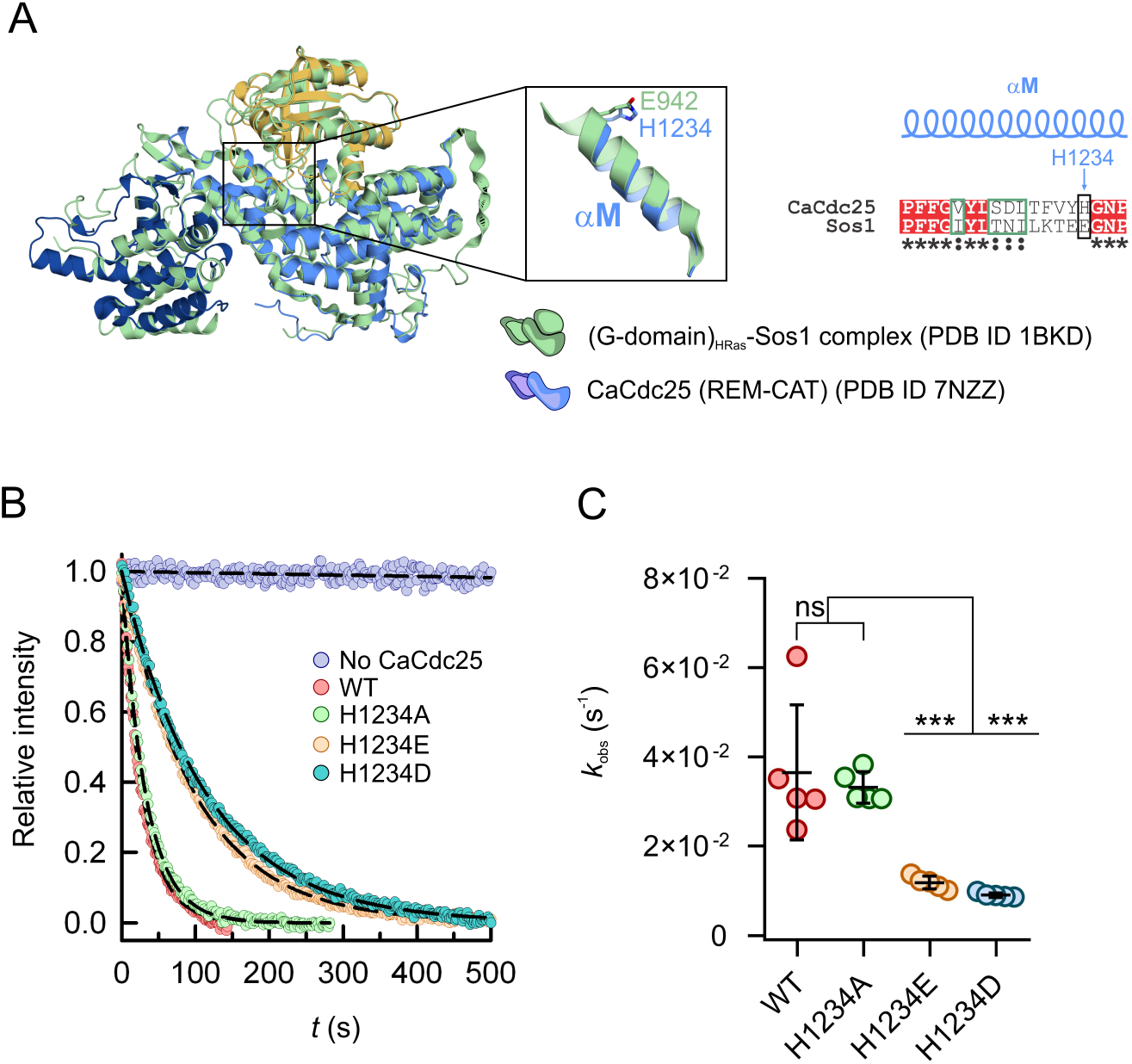
Charge reversal at position 1234, in the unique and conserved in the most common human pathogens α-helical hairpin, diminishes the CaCdc25 nucleotide exchange activity. A) Superposition of the structure of catalytic region of CaCdc25 (PDB code 7NZZ; blue cartoon) with that of the complex of the G-domain of HRas with Sos1 (PDB code 1BKD; grey cartoon). The inset shows the helix of Sos1 essential for Ras nucleotide exchange, aM in CaCdc25, located in the α-helical hairpin, highlighting that the important E942 of Sos1 occupies a position equivalent to that of H1234 in CaCdc25. The sequence alignment of helix aM in *C. albicans* CaCdc25 (UniProtKB entry P43069) with that of human Sos1 (UniProtKB entry Q07889) is shown on the right, with strictly conserved positions in inverted type on a red background, similar positions surrounded by a green rectangle, and H1234 and E942 highlighted with a black rectangle. B) Representative nucleotide exchange reactions of CaRas1 G-domain (at 200 n*M*), isolated or with 100 n*M* of CaCdc25 (wild-type or variants H1234A, H1234E and H1234D). C) Scatter plot of the *k*_obs_ values determined from fitting a single exponential decay model to the nucleotide exchange experimental data shown in B. Five independent experiments were performed. Statistically significant differences (one-way ANOVA and Dunnett’s multiple comparison test) are indicated by *** (p≤0.001) and non-significant by ns (p?0.05).

## Discussion

The extraordinarily flexible adaptation of *C. albicans* to fluctuations in the host microenvironment is a key aspect of its biology, critical for the commensal to pathogen transition of this opportunistic human pathogen (Brown et al., 2014). The *C. albicans* Ras/cyclic AMP (cAMP) signal transduction pathway regulates the expression of several virulence traits involved in host recognition, tissue invasion and evasion of host defense mechanisms. Disrupting the activation of CaRas1, which triggers the signaling pathways that drive the virulence-linked morphological transition from yeast form to a hypha state, is therefore a promising approach for the development of anti-fungal therapies. However, the highly conserved G-domain of CaRas1 (66% amino acid sequence identity with human HRas, with nearly full conservation on the structural elements P-loop, Switch I, Switch II and NKxD motif (Figure S10)) and the striking conservation of Ras activation mechanisms has so far impeded the rational design of molecules targeting the activation of this central environmental sensor in *C. albicans* and in related pathogenic species (LeBlanc et al., 2020; Pentland et al., 2017). Indeed, only a short N-terminal extension in the Ras protein of the human fungal pathogen *Aspergillus fumigatus*, which is absent in Ras homologs of higher eukaryotes, has been proposed as potential target (Al Abdallah et al., 2016). Here we characterize the biochemical and structural features of CaRas1 and its activator CaCdc25 and uncover structural elements specific to *Candida spp.* that could be potentially exploited for the development of novel anti-fungal therapies.

### A structured hypervariable low complexity C-terminal tail in CaRas1

Downstream the highly conserved G-domain of CaRas1, there is a hypervariable region, unusual among Ras-family proteins, which contains low complexity sequence segments, including PolyQ repeats and a Q/N rich stretch, and is predicted to be intrinsically disordered. Cleavage of membrane-associated CaRas1 at its hypervariable region has been proposed as a mechanism restricting CaRas1-mediated activation of the downstream adenylate cyclase (CaCyr1) and, consequently, hyphal growth (Piispanen et al., 2013). In line with previous studies that demonstrated that the hypervariable region of CaRas1 does not play a direct role in CaCyr1 activation (Piispanen et al., 2013), our data reveal that it also does not influence the interaction with the GEF catalytic region of CaCdc25 or its nucleotide exchange activity. So far, the functional role of the hypervariable region of CaRas1 remains elusive, but the structural characterization here reported can provide new clues to unveiling the function of this unusual C-terminal domain.

CaRas1 displays an elongated rod-like shape, with the highly conserved and globular G-domain localizing to one of the ends. The hypervariable region of CaRas1, unstructured according to several predictors, harbors structured elements that account for the extended conformation of the protein. Up close, the hypervariable region consists of two characteristic segments: one composed of a first polyQ region and a Q/N-rich stretch (residues 178-221) and the other comprising a second Q-rich region (residues 248-267, ~70% Q) upstream of the terminal CCaaX motif. The region connecting these two segments (residues 222-247) has a significant content of disorder-favoring glycine (~30%) and serine/alanine (~20%) residues. The first part of the hypervariable region of CaRas1 harbors α-helical elements, as assessed by CD and in agreement with the estimation of several secondary structure predictors. We propose that the terminal helix of the G-domain extends through the first polyQ region until near the end of the Q/N stretch, in line with the susceptibility of the region after N212 to proteolysis in *C. albicans* cells (Piispanen et al., 2013), and in accordance with the predicted model by AlphaFold2 (Figure S11). A similar, although much shorter, helical extension has been described for the human hypervariable region of KRAS4b, which contains a polybasic region formed by a stretch of lysines (Dharmaiah et al., 2016). Additionally, the second polyQ region of CaRas1 seems to be involved in the formation of a stabilizing coiled-coil structure, in line with the LOGICOIL (Vincent et al., 2013) prediction (Figure S12). Indeed, EOM evaluation of the flexibility of the C-terminal part of the hypervariable region (residues 213-290), fixing the G-domain and the helix extension (until N212) (Figure S11*A*), suggested absence of full flexibility and preference for a compact arrangement (Figure S11*B,C*), possibly through interaction with the N-terminal moiety of the hypervariable region, an arrangement also adopted in one of the AlphaFold2 models generated for the molecule (Figure S11*D,E*).

Interestingly, heterotypic and homotypic interactions involving α-helical and intrinsically disordered regions have been shown to mediate phase separation of septal pore-associated proteins in intracellular compartments required for septal biogenesis and cellular communication (Lai et al., 2012). PolyQ repeat-containing proteins have been frequently associated with protein complex assembly and signaling processes (Chavali et al., 2020; Schaefer et al., 2012) and protein self-assembly (Almeida et al., 2013), with impact on resistance to stress and environmental adaptation in yeast (Chavali et al., 2017; Sampaio et al., 2009; Zhou et al., 2017). Recombinant CaRas1 has an intrinsic propensity to self-assemble in the absence of glycerol (data not shown), suggesting that the membrane-bound hypervariable domain might be involved in self-association processes with influence on the regulation of membrane morphology and hyphal growth. This partially structured region, including polyQ stretches, which has similar amino acid composition in homologous Ras1 proteins from other opportunistic fungal pathogens and is absent in the *S. cerevisiae* and human homologues, given its uniqueness can eventually offer new opportunities for the rational design of novel antifungals.

### CaCdc25: a hyperactive Ras1 nucleotide exchange factor with a pathogen-specific region in the α-helical hairpin

CaRas1 is activated by CaCdc25, triggering the virulence-related MAPK and cAMP signaling pathways. The activated form of CaRas1, CaRas1-GTP, binds to CaCyr1 and induces a cAMP pulse that promotes the yeast-to-hypha transition (Fang and Wang, 2006). Whereas the recognition mode of CaRas1 by the catalytic region of CaCdc25 seems to be similar to that of HRas by Sos1, the three-dimensional structure of the REM-CAT tandem of CaCdc25 revealed poor conservation of the residues located in the CaRas1 interaction surface, in particular in the α-helical hairpin, which is critical for triggering the conformational changes required for GDP release. Therefore, and in contrast to the G-domain of CaRas1, where high conservation across species hinders the structure-based design of pathogen-specific targeting molecules, the observed divergences in CaCdc25 make it a potential target to specifically inhibit CaRas1 activation.

Interestingly, the observed stimulation of CaRas1 nucleotide exchange by CaCdc25 was unusually high, suggesting that this GEF is optimally poised for CaRas1 activation. In fact, the nucleotide exchange activity of CaCdc25 is 5- to 2000-fold higher than that of characterized human GEFs (Carabias et al., 2020; Popovic et al., 2013, 2016) (Table S3). This high level of activity of CaCdc25 could result from the stabilization of the α-helical hairpin in an active conformation, as inferred from its crystallographic structure, or from the unique chemical environment of the hairpin, which could influence nucleotide release and therefore CaRas1 activation. Indeed, the nature of the residue at position 1234 seems be critical for the activity of CaCdc25, since the introduction of charge-reversing substitutions at this position (as the point variants H1234D and H1234E) originated less active variants. It is also noteworthy that both H1234 and a small region of the hairpin are strictly and uniquely conserved in the fungal CTG-clade (Figure S9), which groups the most common human fungal pathogens, suggesting a possible correlation between GEF activity and pathogenicity. All these unique features of the sole Ras1-specific GEF identified so far (Piispanen et al., 2011) highlight the relevance of CaCdc25 for pathogenicity, laying the foundation for designing molecules capable to modulate specifically CaRas1 nucleotide exchange and, consequently, the cAMP and MAPK signaling pathways, which regulate critical virulence factors in *C. albicans*.

## Materials and methods

### Cloning and site-directed mutagenesis

DNA sequences encoding the full-length *C. albicans* CaRas1 (UniProt P0CY32) and the fragment comprising the first 166 amino acids were amplified from a synthetic gene by PCR and cloned into the pETGB1a expression vector using the NcoI and Acc65I restriction sites. The resulting constructs encode an N-terminal Gb1-tag, 6×His-tag and TEV cleavage site upstream the CaRas1 protein sequences. The CaRas1 fragment 1-213 (CaRas1-213) expression vector was obtained by inserting a stop codon at position N214 by site directed mutagenesis in the vector encoding the full-length protein. A sequence encoding the fragment comprising amino acids 860-1333 of CaCdc25 (UniProt P43069) was amplified from a synthetic gene by PCR and cloned into the NcoI and Acc65I restriction sites of the derivative of the pETGB1a expression vector (EMBL, Heidelberg, Germany), pETMBP_1. This construct encodes an N-terminal MBP-tag, 6×His-tag and TEV cleavage site upstream the CaCdc25 protein fragment. CaRas1 and CaCdc25 point mutations were introduced by PCR using the QuikChange method. The correctness of all constructs was verified by automated DNA sequencing (oligonucleotides used for gene amplification and site directed mutagenesis are listed in Table S4 and S5, respectively).

### Protein expression and purification

CaRas1 proteins were expressed in *Escherichia coli* BL21 (DE3). Cells were precultured at 37 °C overnight in lysogeny broth (LB) supplemented with 50 mg L^-1^ kanamycin. Twenty mL of the preculture were used to inoculate 2-L flasks containing 500 mL LB, 50 mg L^-1^ kanamycin. When cultures reached an OD600 of 0.6-0.8 after ~1 h shaking at 37 °C, protein production was induced by adding 900 μ*M* isopropyl-β-d-thiogalactopyranoside (IPTG) and expression proceeded for 3 h at 37 °C. Cells were harvested by centrifugation for 30 min at 4 °C and 3,500 *g* and the resulting pellets were resuspended in 20 m*M* Tris-HCl pH 7.5, 500 m*M* NaCl, 20 m*M* imidazole, 5 m*M* β-mercaptoethanol, 5 m*M* MgCl_2_, 5% (*v*/*v*) glycerol (buffer A) and frozen at - 80 °C. Upon thawing the pellet, 1 m*M* PMSF, 0.2 mg mL^-1^ lysozyme, 12.5 m*M* MgCl_2_, and 10 μg mL^-1^ DNAse were added to the cell suspension and stirred for 1h on ice. Next, the cells were disrupted by sonication on ice (5 min at 20% amplitude). The cell lysate was centrifuged for 30 min at 4 °C and 39,200 *g* and the supernatant was loaded onto a 5-mL nickel NTA agarose column (Agarose Bead Technologies) equilibrated with buffer A. Bound protein was eluted with buffer A containing 500 m*M* imidazole. The purity of the eluted fractions was assessed by SDS-PAGE and those containing recombinant protein were pooled. The protein was dialyzed overnight at room temperature against 20 m*M* Tris-HCl pH 7.5, 150 m*M* NaCl, 5 m*M* β-mercaptoethanol, 5 m*M* MgCl_2_, 5% (*v*/*v*) glycerol in the presence of 0.1-0.5 mg His-rTEV to remove the N-terminal MBP-tag and 6×His-tag. Pre- and post-digestion samples were analyzed by SDS-PAGE to confirm the cleavage. The His-rTEV and traces of undigested protein were removed with a second immobilized metal affinity chromatography step, in the same conditions as described above for the first. Target proteins were collected in the flow-through. The protein solution was concentrated to 5 mL in a 10 kDa molecular weight cutoff centrifugal ultrafiltration device (Millipore) and loaded onto a Sephacryl S100 HR 26/60 size-exclusion chromatography column (GE Healthcare), with a mobile phase adequate to the downstream use of the protein. Fractions containing pure protein, as assessed by SDS-PAGE, were combined, concentrated by ultrafiltration, flash frozen in liquid N2 and stored at −80 °C, except for crystallization experiments, where the sample was used immediately. Protein concentrations were determined using the extinction coefficient at 280 nm calculated with the ProtParam tool (http://web.expasy.org/protparam/).

The expression and purification procedures for both wild type of the catalytic region of CaCdc25 and variants were as described for CaRas1, except for the expression strain (*Escherichia coli* BL21 (DE3) Rosetta), the antibiotics used (50 mg L^-1^ kanamycin and 30 mg L^-1^ chloramphenicol) and the induction conditions (200 μ*M* IPTG for overnight induction at 20 °C). Additionally, all buffers were the same as for CaRas1, except for the exclusion of 5 m*M* β-mercaptoethanol, 5 m*M* MgCl_2_ and 5% (*v*/*v*) glycerol. Since CaCdc25 precipitates on ice, all procedures with the purified protein were carried out at room temperature.

CaRas1/REM-CAT complexes were prepared by mixing equimolar amounts of each protein in 20 m*M* sodium phosphate pH 7.5, 150 m*M* NaCl, 1m*M* EDTA, 5% (*v*/*v*) glycerol, 3 m*M* DTT (buffer B). The mixture was incubated for 3 h at room temperature and loaded onto a Superdex 200 10/300 GL (GE Healthcare) size-exclusion chromatography column using buffer B as mobile phase. Fractions containing both proteins, as assessed by SDS-PAGE, were pooled.

### Crystallization

CaCdc25 crystals were obtained at 20 °C by vapor diffusion from sitting drops composed of equal volumes (1 μL) of protein solution (7.2 mg mL^-1^ in 10 m*M* Tris-HCl pH 7.5, 100 m*M* NaCl) and precipitant (23% (w/v) PEG 3350, 0.25 *M* sodium malonate dibasic monohydrate). Prior to data collection the crystals were cryoprotected in crystallization solution supplemented with 25% (*v*/*v*) glycerol and flashed-cooled in liquid N2.

### Data collection and processing

A low resolution (~3 Å) diffraction dataset was collected from two isomorphous crystals of the catalytic region of CaCdc25 (dataset A, Table S2; https://doi.org/10.15785/SBGRID/860 and https://doi.org/10.15785/SBGRID/861) and used for initial structure solution. A partial model derived from dataset A was used as template for molecular replacement (see Structure solution and refinement, below) in a higher resolution (2.45 Å) dataset (dataset B, Table S2; https://doi.org/10.15785/SBGRID/859), which allowed building and refinement of the final model of CaCdc25. All diffraction data were collected at 100 K on the BL13-XALOC beamline (Juanhuix et al., 2014) of the ALBACELLS synchrotron (Cerdanyola del Vallès, Spain). Diffraction data were processed with XDS (Kabsch, 2010a), Pointless (Evans, 2006), and Aimless (Evans and Murshudov, 2013) as implemented in the autoPROC pipeline (Vonrhein et al., 2011). Data sets were scaled and merged with XSCALE (Kabsch, 2010b). All crystals belong to the orthorhombic space group *P*2_1_2_1_2_1_ (Table S2) and contain two monomers of CaCdc25 in the asymmetric unit.

### Structure solution and refinement

The structure of catalytic region of CaCdc25 was solved using dataset A by molecular replacement (MR) with Phaser (McCoy et al., 2007) as implemented in the MrBUMP pipeline (Keegan and Winn, 2008) of the CCP4 suite (Winn et al., 2011). An apparent solution was found using the crystal structure of the REM-CAT region of Sos1 obtained in complex with HRas and ligands (PDB entry 4US0) (Winter et al., 2015) but after some rounds of automatic restrained refinement with REFMAC (Murshudov et al., 2011) the value of *R*free was stuck at >0.50. In order to generate other templates for MR, the initial solution of Phaser was subjected to normal mode analysis using the elNémo server (https://www.sciences.univ-nantes.fr/elnemo/) (Suhre and Sanejouand, 2004). Eleven conformations that represent global motions corresponding to the lowest-frequency mode (200 amplitude perturbation in the direction of a single normal mode with a step size of 40) were generated. Each of these conformations was used as MR search model, and although an improved MR solution was obtained with one of these models, refinement could not be improved and the value of *R*free remained above 0.50. The best model was then subjected to smooth deformation with the morph module of Phenix (Terwilliger et al., 2013) and a substantial improvement in the quality of the electron density maps was observed after 2 morphing cycles (6 Å radius of morphing), accompanied by improved statistics (*R*_work_ = 0.38 and *R*_free_ = 0.48). This model was completed with alternating cycles of refinement with Phenix (Adams et al., 2010) and manual model building with Coot (Emsley et al., 2010). It then became evident that the relative positions of the REM and CAT domains are slightly different in the two molecules of CaCdc25 in the asymmetric unit, probably accounting for the difficulties experienced during the phasing stage. The final refined model derived from dataset A (*R*_work_ = 0.31 and *R*_free_ = 0.34) was used as MR search model with dataset B, and further improved with iterative cycles of refinement (with Phenix (Adams et al., 2010)) and model building (with Coot (Emsley et al., 2010)). The final model (*R*_work_ = 0.19 and *R*_free_ = 0.24) includes residues 883-1035 and 1039-1306 of molecule A and residues 883-1034 and 1039-1305 of molecule B, with 98% of the main-chain torsion angles in the favored regions of the Ramachandran plot. Detailed refinement statistics are given in Table S2. All crystallographic software was supported by SBGrid (Morin et al., 2013).

### Isothermal titration calorimetry (ITC)

ITC experiments were carried out at 25 °C using a VP-ITC system (MicroCal, Northampton, MA, USA). Two solutions of 21 or 42 μ*M* CaCdc25 (in 20 m*M* sodium phosphate pH 7.4, 250 m*M* NaCl, 5 m*M* β-mercaptoethanol, 5 m*M* MgCl_2_, 5% (*v*/*v*) glycerol) were titrated with solutions of 482 μ*M* CaRas1-FL and 370 μ*M* CaRas1-166, respectively, in the same buffer as the CaCdc25 sample and incubated with 10 m*M* EDTA for 30 min prior to loading in the injection syringe. Titrations were done with one initial injection of 3 μL followed by 13 sequential injections of 20 μL each, with 15 μcal s^-1^ reference power, 307 rpm stirring speed and 240 seconds spacing. Heat exchange from the first injection was not used in the analysis. Data were analyzed using the Origin 7 software package (MicroCal) and corrected by the heat of injection calculated from the basal heat remaining after saturation and confirmed by titration into buffer only as control. A single-site model was applied to obtain the stoichiometry (*N*), and association constant (*K*_a_) using a nonlinear squares algorithm.

### Circular dichroism measurements

CaRas1 proteins were dissolved in buffer containing 10 m*M* sodium phosphate pH 7.5, 100 m*M* NaF, 0.5 m*M* MgCl_2_, 5% (*v*/*v*) glycerol or in the same buffer with 50% (*v*/*v*) TFE, to a final concentration of 0.1 mg mL^-1^. CD spectra were recorded using a Jasco J-815 CD spectrometer in a 0.1 cm quartz cuvette for far-UV CD spectroscopy, in a spectral range of 190 nm to 240 nm.

### In vitro nucleotide exchange activity assay

Guanine nucleotide exchange activity was measured by following changes in the fluorescence of the GDP derivative mant-dGDP [2’-Deoxy-3’-O-(N-Methyl-anthraniloyl) guanosine-5’-diphosphate sodium salt (Jena Bioscience GmbH)] (Margarit et al., 2003; Rehmann, 2006). CaRas1 proteins were loaded with mant-dGDP by incubating a 200 μ*M* solution of protein with 2 m*M* mant-dGDP in 20 m*M* Tris-HCl pH 7.5, 50 m*M* NaCl, 4 m*M* EDTA, 1m*M* DTT for 1.5 h at 4 °C (200 μL final volume). The loading reaction was stopped by addition of 10 m*M* MgCl_2_ (30 min, 4 °C). Precipitate formed by addition of MgCl_2_ was removed by centrifuging the sample at 16,000 *g* (30 min, 4 °C). Excess nucleotide was removed by SEC on a Superdex 200 (10/300) column pre-equilibrated with 20 m*M* Tris-HCl pH 7.5, 50 m*M* NaCl, 10 m*M* MgCl_2_. Protein-containing fractions were pooled and concentrated using centrifugal ultra-filtration devices (10 kDa cut-off; Millipore). Protein concentration was determined using a Bradford assay with a standard curve calibration with BSA. Fluorescence measurements were performed on a Fluoromax-4 spectrofluorometer (Horiba-Jobin Ybon), with excitation at 496 nm (1 nm bandwidth) and emission at 518 nm (10 nm bandwidth). Nucleotide exchange experiments were performed at 25 °C in 50 *mM* Tris-HCl pH 7.5, 150 m*M* NaCl, 5 m*M* MgCl_2_. A typical exchange reaction mix contained 200 n*M* CaRas1-mant-dGDP and the reaction was started by addition of 100 n*M* CaCdc25 and 40 μ*M* unlabeled GDP (200-fold molar excess with respect to CaRas1). The exchange rate constant (*k*_obs_) was determined by fitting a single exponential decay model to the time-dependent decrease of fluorescence intensity. Statistical analysis was performed by one-way analysis of variance (ANOVA) followed by Dunnett’s multiple comparisons test and was done with the program GraphPad Prism 8.

### SAXS measurements and data analysis

SAXS data were measured at the P12 beamline of the European Molecular Biology Laboratory (EMBL) at the Deutsches Elektronen-Synchrotron (Hamburg, Germany) using radiation of 1.24 Å wavelength and a Pilatus 6M detector (Dectris) (Blanchet et al., 2015). Samples were equilibrated in the buffers described in Table S1, concentrated by ultrafiltration on 10 kDa or 30 kDa molecular weight cut-off centrifugal devices (Millipore), and centrifuged at 21,500 g and 4 °C for 30 min to remove any aggregates. Prior to data collection, thawed samples were centrifuged and solutions of various concentrations, in the range of 1.1-31.0 mg mL^-1^, were prepared by 2-fold serial dilutions in order to evaluate the magnitude of interparticle effects. All samples and their corresponding buffers were measured consecutively in standard “batch” mode using an automated sample changer, which ensures continuous flow. Thirty frames (0.1 s exposure) for each protein/buffer sample were collected over a 0.0033 to 0.727 Å^-1^ (*q* = (4π sin*θ*)/*2*, where 2*θ* is the scattering angle) scattering vector, at 10 °C (20 °C for CaCdc25 samples). Data were processed and analyzed with the ATSAS 3.0 package (Manalastas-Cantos et al., 2021). Extrapolation from multiple scattering curves at different concentrations to a zero-concentration curve and Guinier analysis were done with PRIMUS/qt. The radius of gyration (*R*_g_) for CaRas1 samples remained constant for all concentrations, within the experimental error. Although a clear dependence of *R*_g_ with concentration was observed for samples of CaCdc25 and of CaRas1/CaCdc25 complexes, extrapolation to zero concentration process was used to alleviate the possible influence of interparticle interactions. The pair distance distribution function, *P*(*r*), was calculated with GNOM. *Ab initio* shape reconstructions were calculated with DAMMIF; multiple reconstructions were superimposed, averaged, and filtered with DAMAVER. Flexibility assessment of the CaRas1 hypervariable region was analyzed using EOM (Bernadó et al., 2007). The scattering profiles of atomic structures were calculated with CRYSOL and missing sequence regions were modelled with MODELLER (Webb and Sali, 2014).

### Sequence and atomic structures analysis

Evolutionary conservation scores were calculated with ConSurf (https://consurf.tau.ac.il/) (Ashkenazy et al., 2016). Superposition of atomic structures were performed with the program Theseus (Theobald and Steindel, 2012). The molecular figures were created by Pymol, version 1.6.0 (Schrödinger) and the secondary structure of CaCdc25 was assigned with DSSP (Kabsch and Sander, 1983).

## Data availability

The X-ray diffraction images (https://doi.org/10.15785/SBGRID/860 and https://doi.org/10.15785/SBGRID/861 for dataset A and https://doi.org/10.15785/SBGRID/859 for dataset B) were deposited with the Structural Biology Data Grid (Meyer et al., 2016). Coordinates and structure factors were deposited at the Protein Data Bank (PDB) under accession number 7NZZ. SAXS data were deposited at the Small Angle Scattering Biological Data Bank (SASBDB) (Valentini et al., 2015) under codes SASDEM6, SASDEN6, SASDEP6, SASDEQ6 and SASDER6. Other data are available from the corresponding authors upon reasonable request.

## Acknowledgements

We acknowledge the EMBL (DESY, Hamburg, Germany) for the provision of SAXS measuring time (proposal SAXS-1020) and Dr. Andrey Gruzinov and Dr. Stefano Da Vela (BioSAXS beamline P12, PETRA III storage ring) for collecting the data. The X-ray crystallography measurements were performed at beamline BL13-XALOC of the ALBA Synchrotron (Cerdanyola del Vallès, Spain) with support from ALBA staff. The support of the X-ray Crystallography and Biochemical and Biophysical Technologies scientific platforms of i3S (Porto, Portugal) is also acknowledged.

This work was funded by the European Union’s Horizon 2020 Research and Innovation programme under grant agreement ID 952334 “PhasAGE”. J.A.M. also acknowledges FCT-Fundação para a Ciência e a Tecnologia for research contracts (POCI-01-0145-FEDER-029221 and PTDC/BIA-BQM/2494/2020).

## Author contributions

J.A.M., and S.M.-R. designed research; J.A.M., A.C., and Z.S. performed research; J.A.M., J.M.P., P.J.B.P., and S.M.-R. analyzed data; and J.A.M., P.J.B.P., and S.M.-R. wrote the paper.

## Conflict of interest

The authors declare no competing financial interests.

## Notes

### Competing Interest Statement

The authors have declared no competing interest.

