## Supplementary Material for "Pathogen-specific structural features of two key players in *Candida albicans* morphogenetic switch"

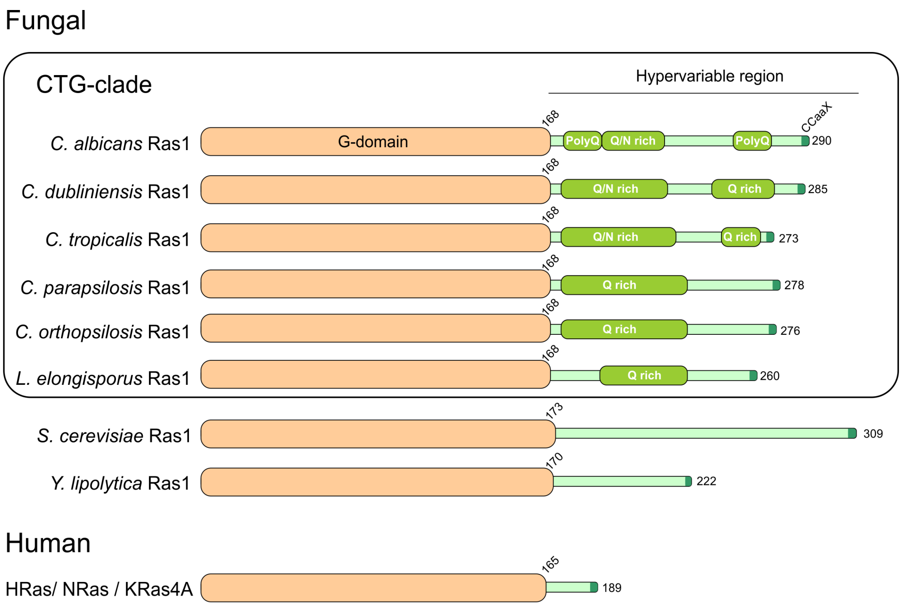


**Figure S1. Domain organization of fungal and human Ras proteins.** A main difference between fungal and human Ras proteins resides on the length of the hypervariable region, much longer in fungi. The presence of polyQ and Q/N rich regions at the hypervariable region in members of the CTG-clade makes Ras1 unique within the Ras family proteins. *C. albicans* Ras1 (UniProtKB entry P43069), *C. dubliniensis* Ras1 (*Candida* Genome Database (CGD) entry Cd36_24270, *C. tropicalis* Ras1 (CGD entry CTRG_02064), *C. parapsilosis* Ras1 (CGD entry CPAR2_407360), *C. orthopsilosis* Ras1 (CGD entry CORT_0C06700), *L. elongisporus* Ras1 (CGD entry LELG_02372), *S. cerevisiae* Ras1 (UniProtKB entry P01119) and *Y. lipolitica* Ras1 (GenBank entry KAG5356685.1), and HRas (UniProtKB entry P01112), NRas (UniProtKB entry P01111) and KRas4A (UniProtKB entry P01116-1) were selected as representative fungal and human Ras proteins, respectively.

A

| Model | Template (organism) | PDB entry | FFAS-score | Seq. identity, % | $Chi^2$ |
|---|---|---|---|---|---|
| 1 | RRas2 (human) | 2ERY | −92.5 | 62 | 2.5 |
| 2 | Rap1b (human) | 3X1Z | −91.2 | 61 | 2.6 |
| 3 | RalA (human) | 6P0I | −92.1 | 51 | 2.9 |
| 4 | Rap1a (human) | 1GUA | −91.2 | 62 | 3.1 |
| 5 | Ras (Choanoflagellata) | 5WDR | −91.9 | 72 | 3.3 |
| 6 | HRas (human) | 6NTC | −91.7 | 66 | 3.4 |
| 7 | HRas (human) | 5VBE | −91.6 | 66 | 3.5 |
| 8 | Rap2A (human) | 1KAO | −93.1 | 52 | 4.0 |
| 9 | Rheb (human) | 6BSX | −93.0 | 40 | 7.3 |
| 10 | Rheb (Mus musculus) | 4O25 | −92.7 | 40 | 7.9 |
| 11 | KRas G12C (human) | 4L8G | −91.3 | 65 | 8.7 |
| 12 | NRas (Homo sapiens) | 3CON | −90.1 | 68 | 9.6 |

C

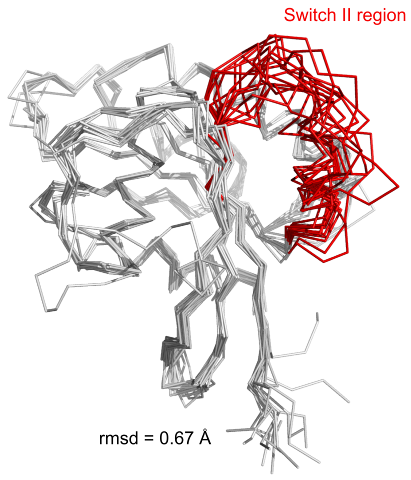


B

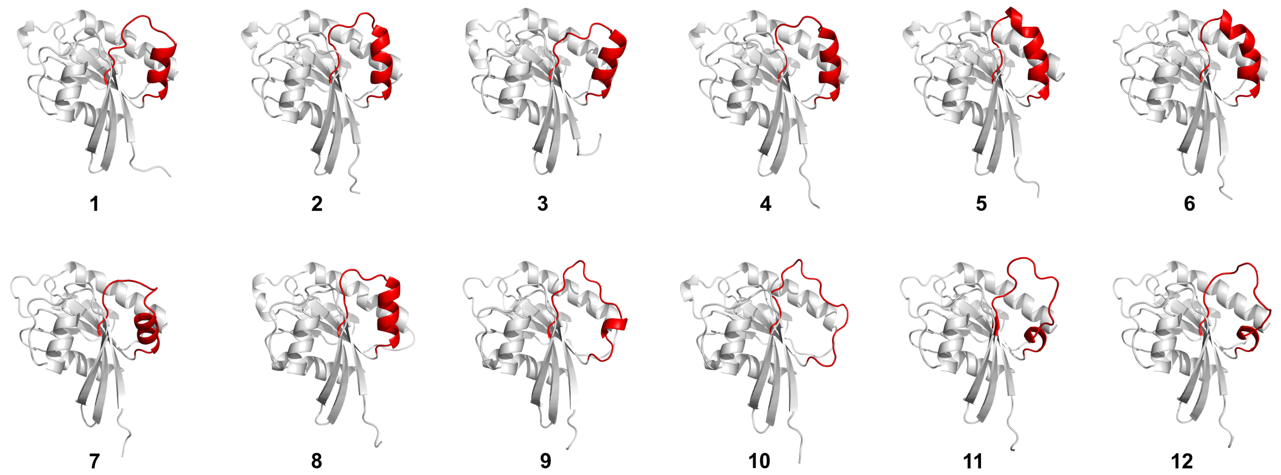


D

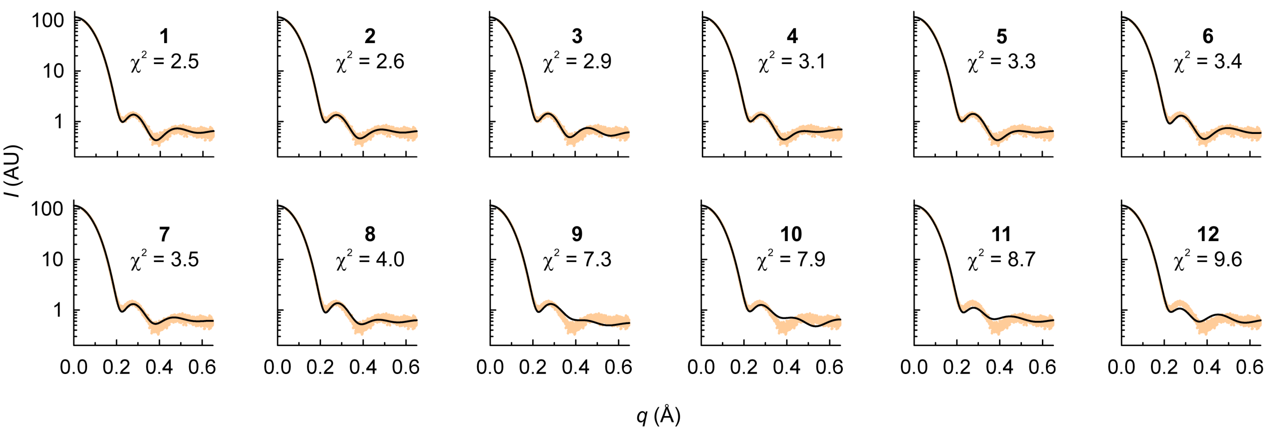


**Figure S2. Structure of the CaRas1 G-domain.** A) Models for the *C. albicans* CaRas1 G-domain using as templates twelve structural homologues (small GTPase proteins with 40-66% amino acid sequence identity) identified with the Fold and Function Assignment System (FFAS03) (Jaroszewski et al., 2011). The models are sorted according to their fit to the SAXS experimental data (as judged by the value of $chi^2$). Despite the low FFAS-score for all models (scores below −9.5 correspond to high-confidence predictions with less than 3% false positives) and the high degree of amino acid sequence conservation, there are notable differences in the SAXS $chi^2$ values, which represent the discrepancy between the theoretical and experimental SAXS curves (see panel D). B) Cartoon representation of the models from panel A. Structural differences are mostly located on the switch II region (colored red). C) Structural superposition of the 12 models (represented as Cα traces), highlighting the variations in the switch II region (colored as in panel B). D) Experimental SAXS scattering curve for the CaRas1 G-domain (orange dots) and theoretical curves estimated from the 12 different models (black line). Models with a switch II region displaying α-helical secondary structure (1-8) originate theoretical scattering curves that better fit the experimental data.

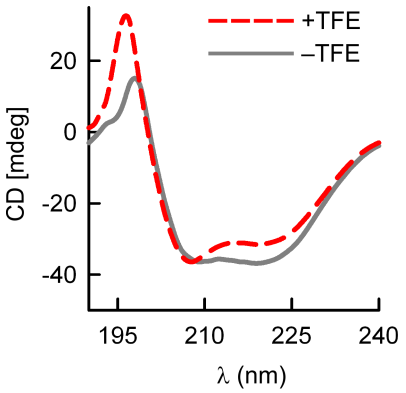


**Figure S3. Inversion of the 220 nm/209 nm ratio in the CD spectrum of CaRas1-FL in the presence of TFE.** Circular dichroism spectra of CaRas1-FL in absence (solid gray line) and in the presence of 50% TFE (dashed red line). A shift in the 220 nm/209 nm ratio from 1.00 to 0.88 was observed when CaRas1-FL was measured in presence of 50% TFE.

A

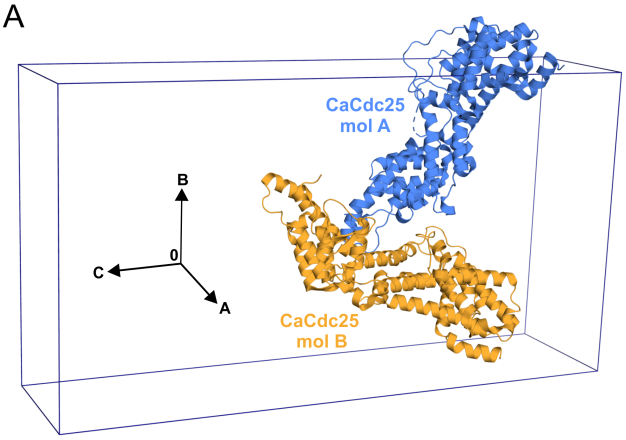


B

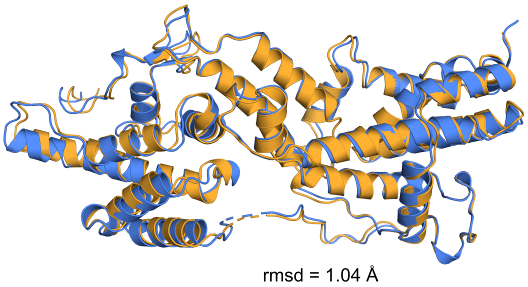


C

| PDB entry | Protein | *Z*-score | % seq. identity | Ref. |
|---|---|---|---|---|
| 5CM8 | Rlf (GEF for Ral) | 31.6 | 17 | (3) |
| 4L9M | RasGRP1 (GEF for Ras) | 27.3 | 18 | (4) |
| 4F7Z | EPAC2 (GEF for Rap1b) | 26.2 | 22 | (5) |
| 3KSY | Sos1 (GEF for Ras) | 23.1 | 25 | (6) |

D

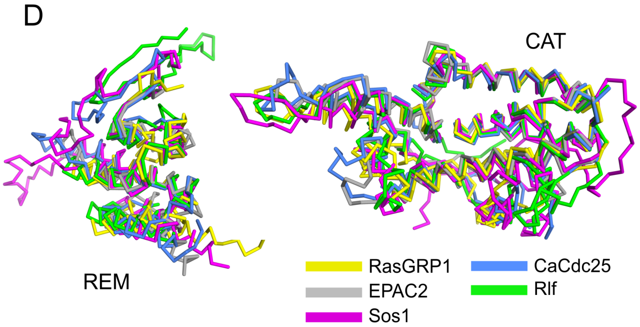


**Figure S4. Overall view of the structures of the two CaCdc25 monomers in the crystallographic asymmetric unit and comparison with other GEF homologues.** A) Cartoon representation of the two molecules (blue: molecule A; orange: molecule B) found in the asymmetric unit (AU) of the catalytic region of CaCdc25. B) Structural superposition of the two CaCdc25 monomers found in the crystallographic AU (colored as in panel A). Slight differences were found in the relative orientation of the CAT and REM domains in the two molecules, probably due to the distinct intermolecular contacts in the crystal packing. C) Structural homologues of CaCdc25 identified with the DALI (Holm, 2020) server using molecule A as query structure. There were only four hits with Z-score > 16, all displaying surprisingly low amino acid sequence conservation (≤ 25% sequence identity), corresponding to PDB entries 5CM8 (Popovic et al., 2016), 4L9M (Iwig et al., 2013), 4F7Z (White et al., 2012) and 3KSY (Gureasko et al., 2010). D) Superposition (Cα traces) of the individual structures of the REM domain (left) and CAT domain (right) of CaCdc25 and the four structural homologues listed in panel C.

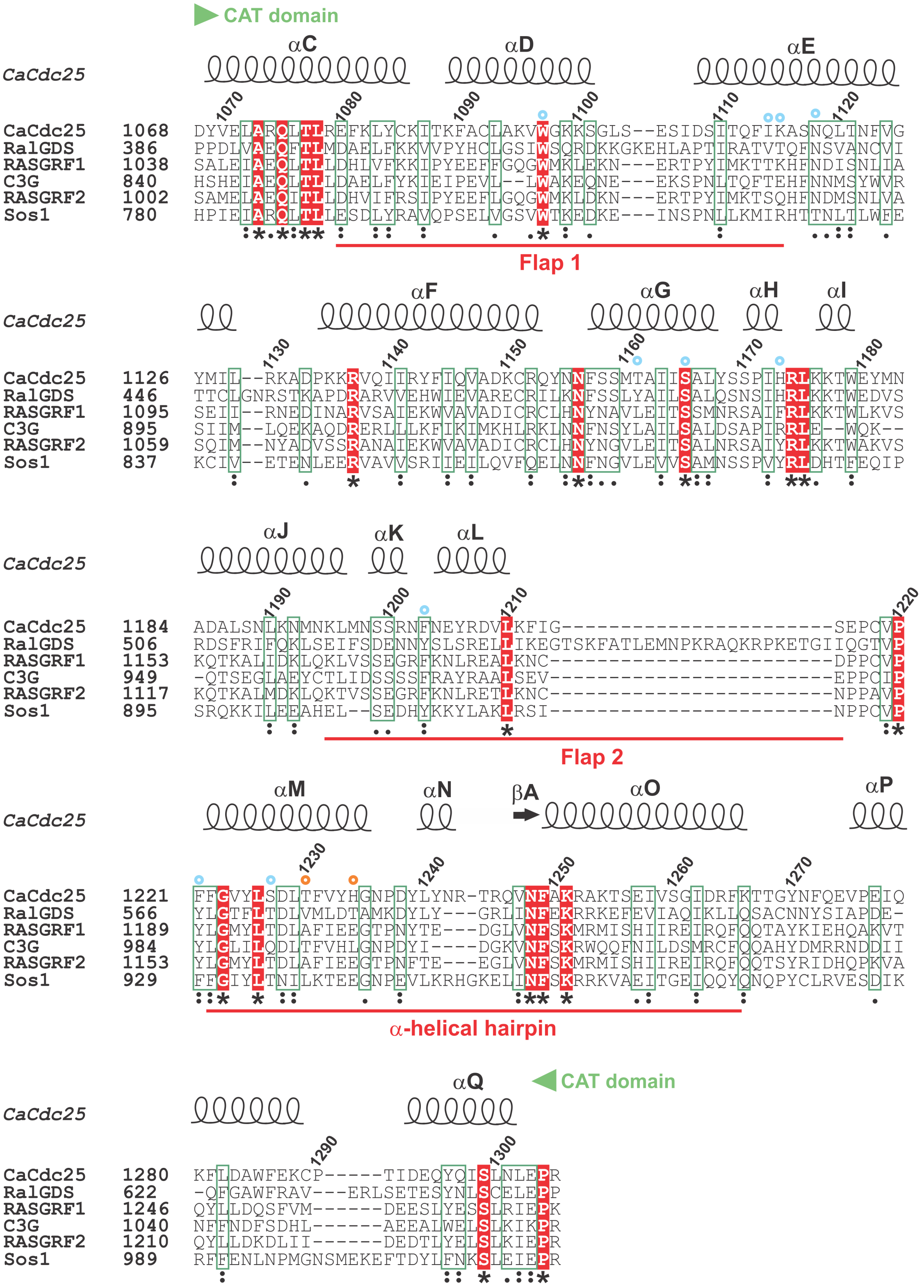


**Figure S5. Multiple amino acid sequence alignment of the CAT domain of CaCdc25 with human homologues.** The amino acid sequence of the CAT domain of *Candida albicans* CaCdc25 (UniProtKB entry P43069) was aligned with those of humans: RalGDS (UniProtKB entry Q12967), RASGRF1 (UniProtKB entry Q13972), C3G (UniProtKB entry Q13905), RASGRF2 (UniProtKB entry O14827) and Sos1 (UniProtKB entry Q07889). Strictly conserved alignment positions are shown in inverted type on a red background. Secondary structure elements for the CAT domain of CaCdc25 are represented above the alignment. The flap 1 and flap 2 regions, as well as the α-helical hairpin are labeled in red. Blue circles indicate Sos1 residues that directly contact HRas in the crystallographic structure of the complex (PDB entry 1BKD) (Boriack-Sjodin et al., 1998), and orange circles denote residues from the nucleotide binding site.

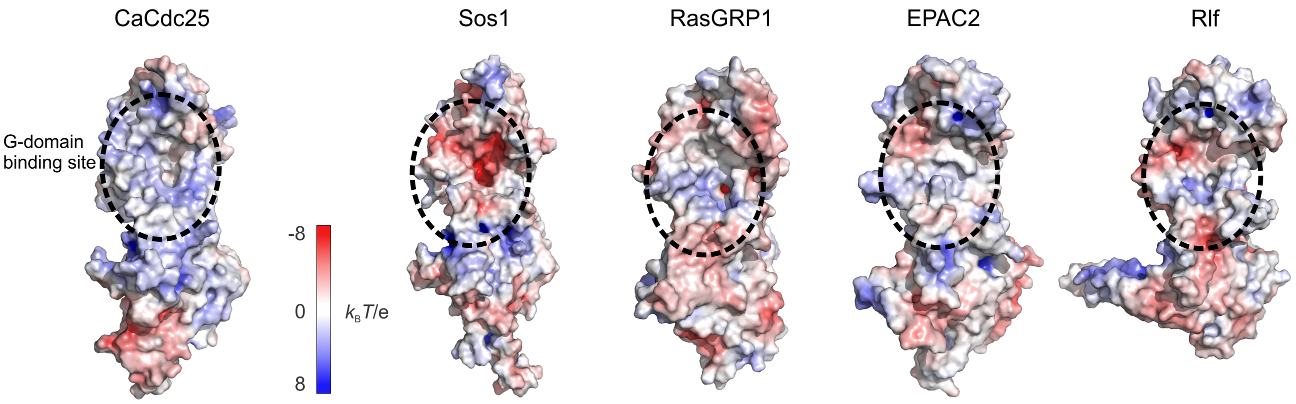


**Figure S6. Comparison of the surface electrostatic potential of *C. albicans* Cdc25-GEF and mammalian structural homologues.** Representation of the molecular surfaces of the CAT and REM domains colored according to their electrostatic potential. Structures of the mammalian homologues Sos1 (PDB entry 3KSY (Gureasko et al., 2010)), RasGRP1 (PDB entry 4L9M (Iwig et al., 2013)) EPAC2 (PDB entry 4F7Z (White et al., 2012)) and Rlf (PDB entry 5CM8 (Popovic et al., 2016)) are represented. The G-domain binding site of each GEF for the corresponding GTPase is indicated by a black dashed circle.

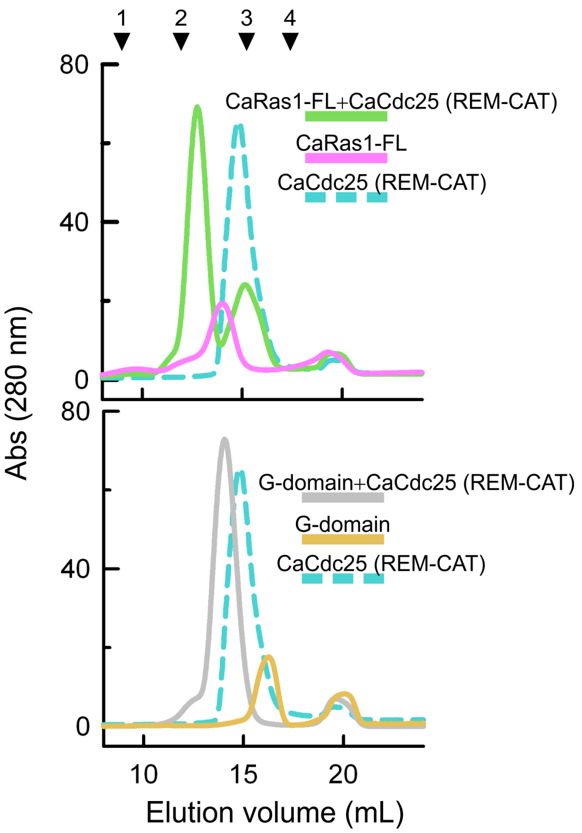


**Figure S7. Analysis by size exclusion chromatography of the complex formed by the catalytic region of CaCdc25 (REM-CAT) and CaRas1-FL or CaRas1 G-domain.** CaRas1/REM-CAT complexes were prepared by mixing equimolar amounts of each protein in 20 m*M* sodium phosphate pH 7.5, 150 m*M* NaCl, 1m*M* EDTA, 5% (*v*/*v*) glycerol, 3 m*M* DTT. The mixture was incubated for 3 h at room temperature and loaded onto a Superdex 200 10/300 GL (GE Healthcare) size-exclusion chromatography column using the above buffer as mobile phase. Chromatograms of the isolated proteins (REM-CAT (blue), CaRas1-FL (pink) and CaRas1 G-domain (orange)) and of the equimolar mixtures of the catalytic region of CaCdc25 with CaRas1-FL (green) and CaRas1 G-domain (grey) are shown. The inverted triangles above the chromatograms mark the position of the elution peaks of the proteins used as standards: (1) thyroglobulin (670 kDa), (2) γ-globulin (158 kDa), (3) ovalbumin (44 kDa), and (4) myoglobin (17 kDa).

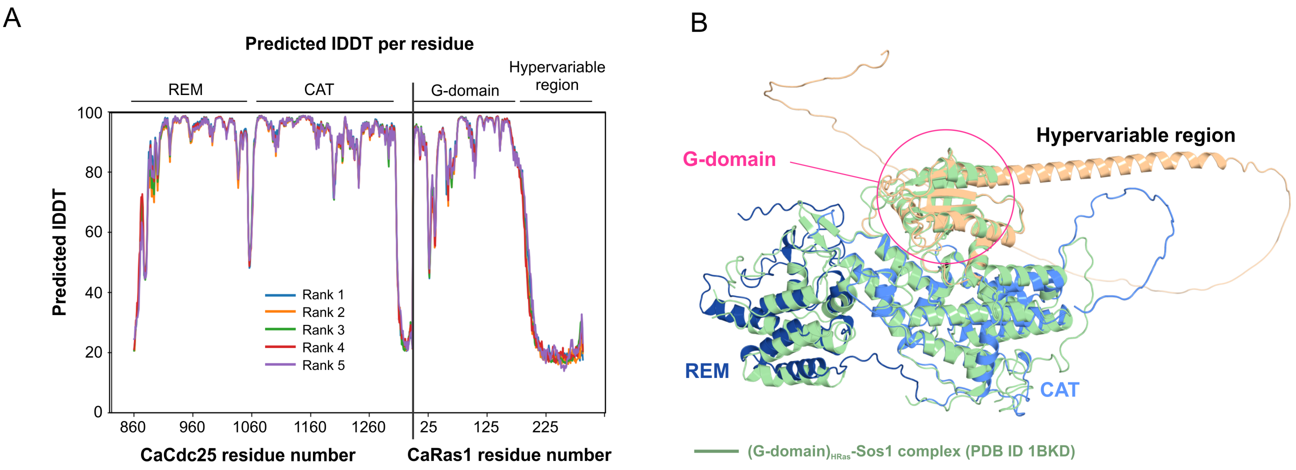


**Figure S8. Predicted structure of the catalytic region of CaCdc25 (REM-CAT domains) in complex with full-length CaRas1 (CaRas1-FL).** A) Per-residue local Distance Difference Test (lDDT) of five predicted structures of CaRas1-FL/REM-CAT complex. The interaction mode of the tandem REM-CAT of CaCdc25 with CaRas1-FL was identical among the five different predictions. Complex predictions were performed with AlphaFold2 using Many-against-Many sequence searching (MMseqs2) through the ColabFold notebook (Mirdita et al., 2022). B) One of the five predicted models by AlphaFold2 for CaCdc25 (REM and CAT domains in dark and light blue, respectively) in complex with CaRas1-FL (orange). The crystallographic structure of the complex of Sos1 with HRas (PDB entry 1BKD (Boriack-Sjodin et al., 1998); colored as palegreen) is superposed to the predicted complex of *Candida albicans* where the position of corresponding G-domains is highlighted by a circle.

A

B

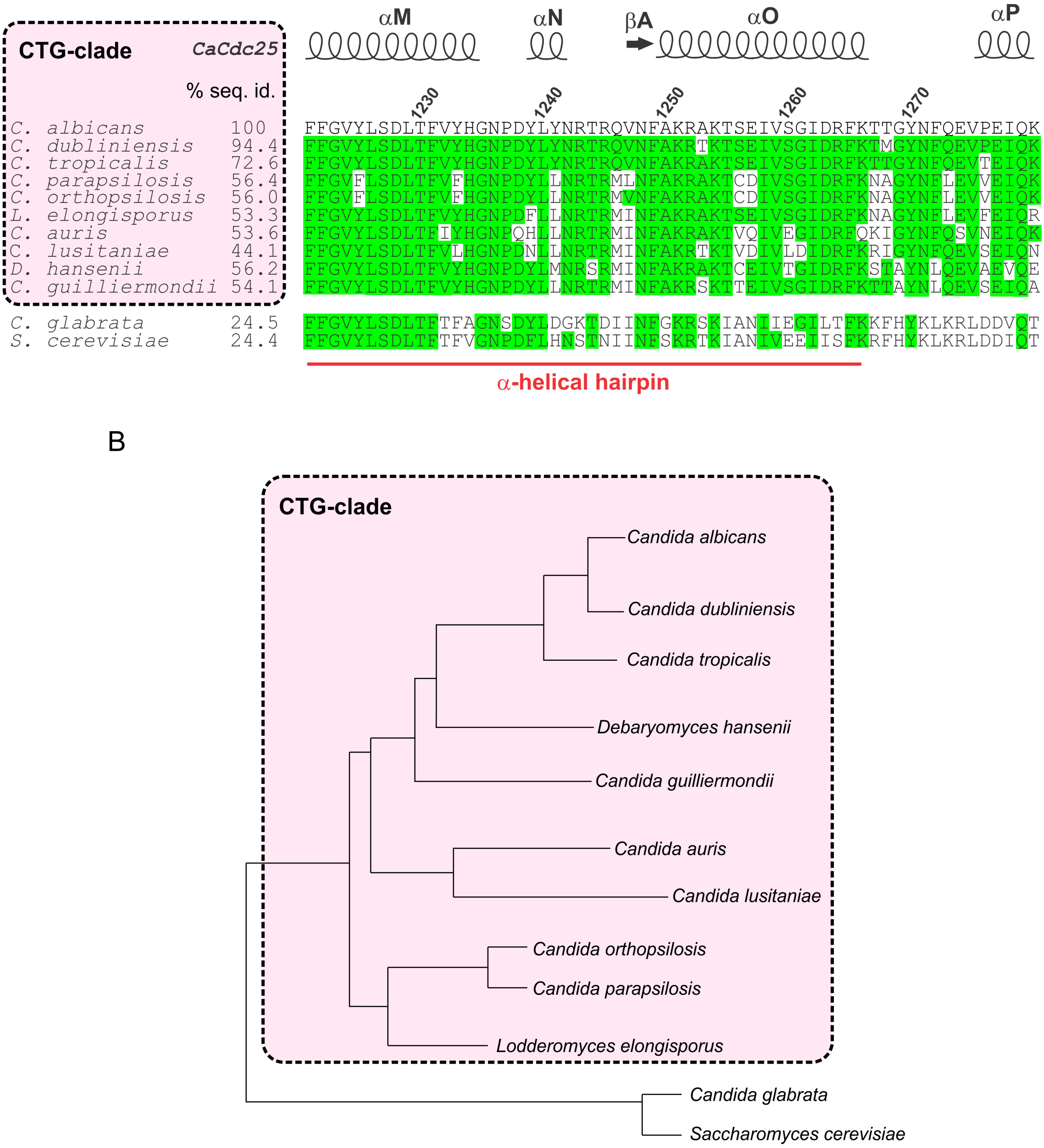


**Figure S9. Helix αM is conserved in common pathogenic fungi.** A) Amino acid sequence alignment highlighting the exclusive conservation of helix αM in only ten identified CaCdc25 homologues from other common human pathogenic fungi. B) Phylogenetic tree of CaCdc25 homologues obtained from the *Candida* Genome Database (http://www.candidagenome.org/) and built with SEMPHY (Ninio et al., 2007).

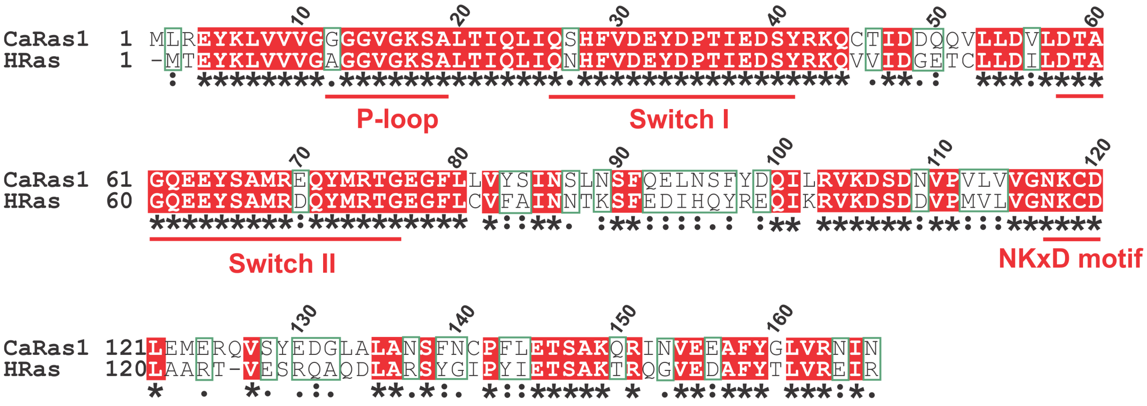


**Figure S10. Amino acid sequence alignment of the G-domain of CaRas1 with that of human HRas.** The amino acid sequence of the G-domain of *Candida albicans* CaRas1 (UniProtKB entry P0CY32) was aligned with that of human hRas (UniProtKB entry P01112). Strictly conserved alignment positions are shown in inverted type on a red background. The P-loop, Switches I and II, and the NKxD motif are labeled in red.

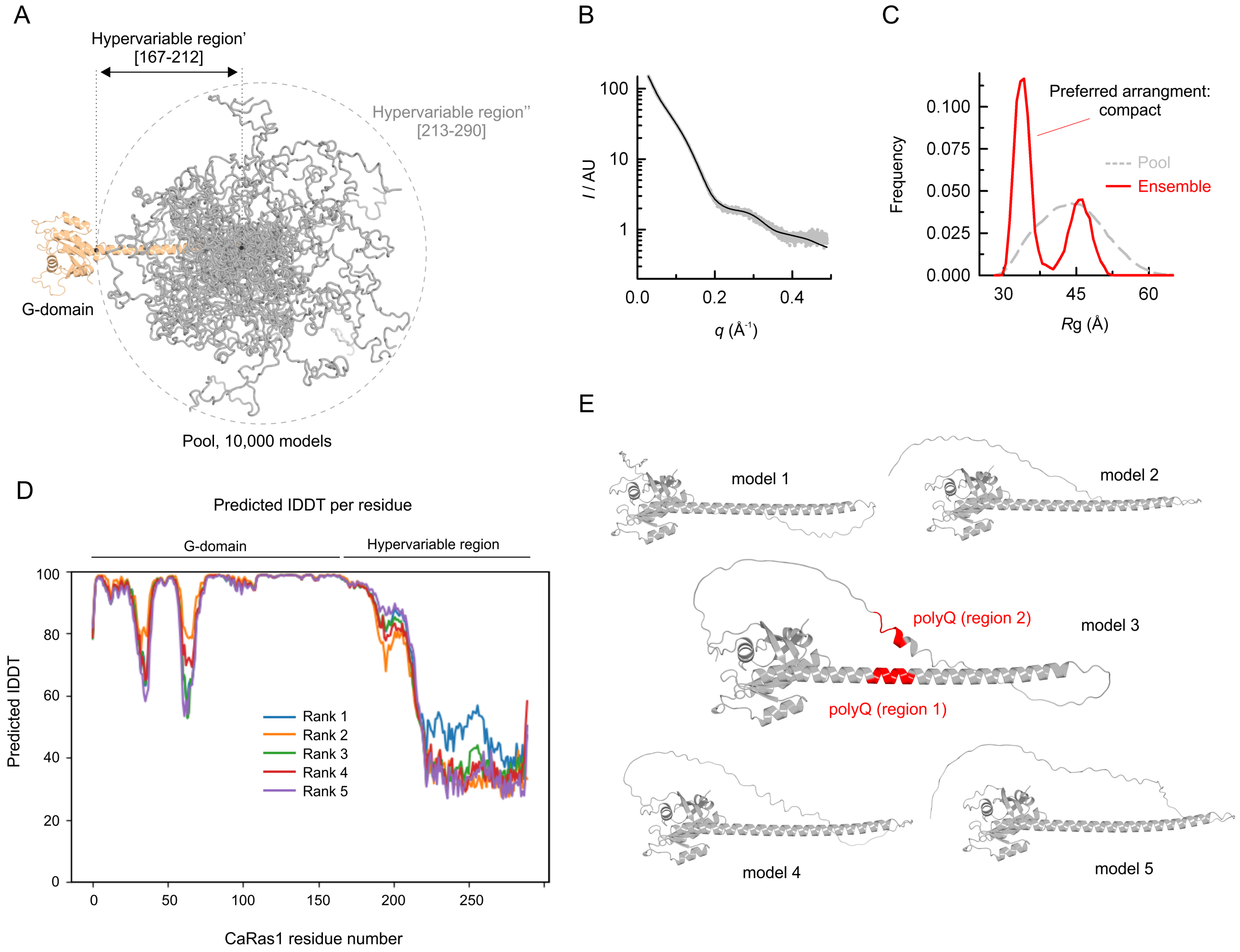


**Figure S11. Flexibility analysis of the segment 213-290 in the hypervariable region of CaRas1 in the full-length CaRas1 structures predicted by AlphaFold2.** A) Representation of the generated pool to evaluate the flexibility of the CaRas1 region 213-290 (colored in gray) by EOM. The pool consisted of 10,000 models, in which both the G-domain and the region 167-212 (colored in orange) were fixed, whereas the part of the hypervariable region, that comprises the residues 213-290, was considered fully flexible (for clarity only 50 models are represented). B) SAXS experimental data of the full-length CaRas1 (represented as gray dots) and the scattering profile calculated for the selected EOM ensemble (represented as a solid black line) which fits the data with a $\chi^2$ of 2.218. C) EOM analysis of the flexibility of the 213-290 region (hypervariable region''). Frequency of radius of gyration (*R*g) distributions in a pool of 10,000 models (gray dashed line) with random orientations of the hypervariable region'', and in the selected ensemble that fits the SAXS data of CaRas1-FL that is shown in panel B (red line). Default parameters were employed using native-like models, allowing constant subtraction (0.149) and curve repetition (both the minimum number of curves per ensemble as well as the number of obtained representative structures, was five). The values for *R*flex(random)/*R*sigma of ~ 73.56% (~ 89.13%) / 0.91 indicate a limited flexibility (number of representative structures, five). D) Per-residue local Distance Difference Test (lDDT) of five predicted structures of full-length CaRas1. Predictions were performed with AlphaFold2 using MMseqs2 through the ColabFold notebook (Mirdita et al., 2022). E) The five structures predicted for full-length CaRas1. In model 3 the two polyQ regions are colored in red.

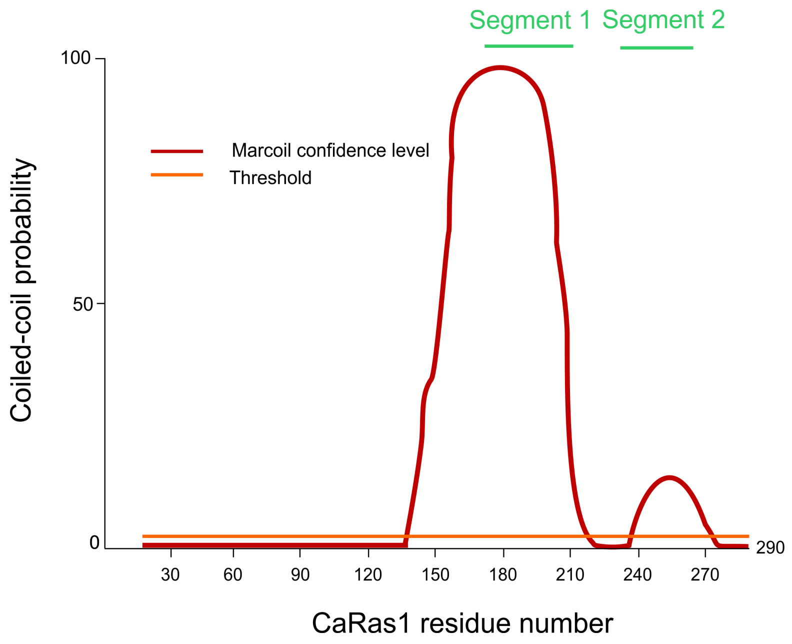


**Figure S12. Prediction of a coiled-coil in CaRas1.** Two segments within the hypervariable region of CaRas1 are predicted by LOGICOIL (Vincent et al., 2013) to form coiled-coil structures.

**Table S1. Small angle X-ray scattering results for CaCd25 (tandem REM-CAT), CaRas1 constructs and CaRas1/REM-CAT complexes[a].**

| **Sample details** | **CaCdc25 (REM-CAT)** | **CaRas1 G-domain** | **CaRas1-FL** | **CaRas1 G-domain/REM-CAT** | **CaRas1-FL/REM-CAT** |
|---|---|---|---|---|---|
| Organism | *Candida albicans* | | | | |
| Source | Expressed in *E. coli* BL21 (DE3) Rosetta | Expressed in *E. coli* BL21 (DE3) | | As described for the individual components | |
| UniProtKB entry (residues in construct) | P43069 (860-1333) | P0CY32 (1-166) | P0CY32 (1-290) | As described for the individual components | |
| $\bar{v}$ from chemical composition ($cm^3$ $g^{-1}$)[b] | 0.738 | 0.733 | 0.719 | 0.737 | 0.731 |
| Particle contrast from sequence and solvent constituents, $\Delta\bar{\rho}$ ($\rho_{protein}$ - $\rho_{solvent}$; $10^{10}$ $cm^{-2}$)[b] | 2.70 (12.29 – 9.59) | 2.77 (12.35 – 9.58) | 3.00 (12.58 –9.58) | 2.72 (12.31 – 9.59) | 2.81 (12.40 – 9.59) |
| Concentration range (mg $mL^{-1}$) | 1.4 – 11.1 | 1.9 – 31.0 | 2.1 – 17.0 | 1.1 – 8.5 | 1.4 – 11.3 |
| *M* from chemical composition (kDa) | 55 | 19 | 33 | 74 | 87 |
| Solvent (solvent blanks taken from SEC flow-through prior to elution of protein) | 20 m*M* sodium phosphate pH 7.5, 150 m*M* NaCl, 5% (v/v) glycerol, 3 m*M* DTT | 20 m*M* Tris-HCl pH 7.5, 150 m*M* NaCl, 5% (v/v) glycerol, 3 m*M* DTT, 5 m*M* $MgCl_2$ | | 20 m*M* sodium phosphate pH 7.5, 150 m*M* NaCl, 1m*M* EDTA, 5% (v/v) glycerol, 3 m*M* DTT | |
| **Structural parameters** | | | | | |
| Guinier analysis | | | | | |
| *I*(0)/c ($10^{-2}$ $cm^2$ $mg^{-1}$)[c] | 3.21 ± 0.01 | 1.01 ± 0.01 | 1.89 ± 0.01 | 4.13 ± 0.01 | 6.15 ± 0.01 |
| $R_g$ (Å) | 31.7 ± 0.1 | 16.1 ± 0.1 | 36.2 ± 0.1 | 32.0 ± 0.6 | 39.7 ± 0.1 |
| $q_{min}$ ($Å^{-1}$) | 0.012 | 0.042 | 0.022 | 0.029 | 0.014 |
| $qR_g$ max | 1.30 | 1.30 | 1.30 | 1.30 | 1.18 |
| Correlation coefficient, $R^2$ | 0.9991 | 0.9995 | 0.9928 | 0.9961 | 0.9976 |
| *M* from bayesian inference [Credibility interval probability] (ratio to predicted) | 58 [91%] (1.1) | 17 [93%] (0.9) | 37 [93%] (1.1) | 53 [93%] (0.7) | 94 [91%] (1.1) |
| *M* from *I*(0)/c (ratio to predicted)[d] | 43 (0.8) | 11 (0.6) | 21 (0.6) | 54 (0.7) | 77 (0.9) |
| *M* from BSA (ratio to predicted) | 43 (0.8) | 14 (0.7) | 25 (0.8) | 55 (0.7) | 85 (1.0) |
| *P*(*r*) analysis | | | | | |
| *I*(0)/c ($10^{-2}$ $cm^2$ $mg^{-1}$)[c] | 3.25 ± 0.01 | 1.02 ± 0.01 | 1.99 ± 0.01 | 4.17 ± 0.01 | 6.28 ± 0.01 |
| $R_g$ (Å) | 32.9 ± 0.1 | 16.0 ± 0.1 | 41.1 ± 0.1 | 33.1 ± 0.6 | 42.8 ± 0.1 |
| $d_{max}$ (Å) | 118 | 41 | 151 | 118 | 177 |
| *q* range ($Å^{-1}$) | 0.012 – 0.300 | 0.042 – 0.497 | 0.020 – 0.34 | 0.021 – 0.25 | 0.014 – 0.20 |
| Total estimate from *GNOM* | 0.73 | 0.62 | 0.59 | 0.82 | 0.71 |
| *M* from *I*(0) (ratio to predicted value)[d] | 44 (0.8) | 11 (0.6) | 22 (0.7) | 55 (0.7) | 79 (0.9) |
| Porod volume ($Å^3$) (ratio $V_P$ / *M* from sequence) | 81,103.9 (1.5) | 26,029.1 (1.4) | 55,522.3 (1.7) | 99,438.5 (1.3) | 148,691 (1.7) |

**Table S1.** Continuation.

| **Shape model-fitting results** | **CaCdc25 (REM-CAT)** | **G-domain** | **CaRas1-FL** | **CaRas1 G-domain/REM-CAT** | **CaRas1-FL/REM-CAT** |
|---|---|---|---|---|---|
| DAMMIF (interactive mode, 15 calculations) | | | | | |
| $q$ range for fitting (Å$^{-1}$) | 0.012 – 0.300 | 0.042 – 0.497 | 0.020 – 0.34 | 0.021 – 0.25 | 0.014 – 0.20 |
| Symmetry, anisotropy assumptions | *P*1, prolate | *P*1, unknown | *P*1, prolate | *P*1, prolate | *P*1, prolate |
| NSD (standard deviation), No. of clusters | 1.05 (0.08), 1 | 0.83 (0.04), 1 | 0.75 (0.05), 1 | 0.89 (0.12), 1 | 0.93 (0.06), 1 |
| $\chi^2$ range | 1.49 – 1.53 | 3.57 – 3.62 | 1.93 – 2.00 | 1.69 – 1.72 | 1.49 – 1.55 |
| Constant adjustment to intensities | 0.235 | Skipped, unable to determine | Skipped, unable to determine | 2.06 | 1.47 |
| Resolution (from SASRES) (Å) | 42 ± 3 | 24 ± 2 | 29 ± 2 | 39 ± 3 | 47 ± 4 |
| $M$ estimate as 0.5 × volume of models (kDa) (ratio to expected) | 52 (0.9) | 10 (0.5) | 29 (0.9) | 65 (0.9) | 98 (1.1) |
| **Atomistic modelling** | | | | | |
| Crystal structures | 7NZZ (This work) | Structure modelled using 2ERY as template | | Structure modelled using 1BKD as template | |
| q range for all modelling | 0.0098 – 0.50 | 0.0395 – 0.65 | | 0.0243 – 0.30 | |
| *CRYSOL* (max. order of harmonics 50) | | | | | |
| Constant subtraction allowed | | | | | |
| $\chi^2$ | 2.86 | 2.71 | | 6.85 | |
| Predicted $R_g$ (Å) | 31.8 | 16.1 | | 30.6 | |
| Vol (Å), Ra (Å), Dro (e Å$^{-3}$) | 72,996 / 1.80/ 0.048 | 24,269 / 1.40 / 0.005 | | 89,971 / 1.80 / 0.003 | |
| No constant subtraction | | | | | |
| $\chi^2$ | 2.92 | 4.49 | | 23.49 | |
| Predicted $R_g$ (Å) | 31.8 | 16.3 | | 30.5 | |
| Vol (Å), Ra (Å), Dro (e Å$^{-3}$) | 71,958 / 1.74 / 0.052 | 22,738 / 1.44 / 0.015 | | 99710 / 1.80 / 0.000 | |
| *EOM* (default parameters, 10 000 models in initial ensemble, native-like models, constant subtraction allowed) | | | | | |
| Minimum number of curves per ensemble | | | 5 | | 5 |
| Curve repetition in the ensemble allowed? | | | Yes | | Yes |
| $\chi^2$ | | | 2.49 | | 2.55 |
| *R*flex(random)/*R*sigma | | | ~73.3% (~84.1%)/0.61 | | ~74.6% (~88.3%)/0.55 |
| Constant subtraction | | | 0.182 | | 0.424 |
| No. of representative structures | | | 5 | | 4 |
| **SASBDB code**[e] | SASDM75 | SASDM55 | SASDM65 | SASDM85 | SASDM95 |

**Table S1.** Continuation.

[a]Description of the accuracy and confidence in the SAXS data and modelling outputs are reported following the *2017 publication guidelines and recommendations for solution small-angle scattering data* (Trewhella et al., 2017). [b]Partial specific volumes, $\bar{v}$, and the particles contrast, $\Delta\bar{\rho}$, were calculated with MULCh (Whitten et al., 2008). [c]Absolute intensities were determined using water as secondary standard. [d]$M$ was calculated as $[N_A\ I(0)/c]/\Delta\rho_M^2$, where $I(0)/c$ is the forward scattering normalized against concentration, $\Delta\rho_M = [\rho_{M,prot} - (\rho_{solv}\ \bar{v})]r_o$ is the scattering contrast per mass, $N_A = 6.023 \times 10^{23}$ mol$^{-1}$ is the Avogadro number, $\rho_{M,prot} = 3.22 \times 10^{23}$ e g$^{-1}$ is the number of electrons per mass of dry protein, $\rho_{solv} = 3.34 \times 10^{23}$ e cm$^{-3}$ is the number of electrons per volume of the aqueous solvent, $\bar{v}$ is the partial specific volume of the protein and $r_o = 2.8179 \times 10^{-13}$ cm is the scattering length of an electron (Feigin and Svergun, 1987; Mylonas and Svergun, 2007; Orthaber et al., 2000). [e]SASBDB, Small Angle Scattering Biological Data Bank (Valentini et al., 2015).

**Table S2. Crystallographic data collection and refinement statistics**

| | CaRas1 guanine-nucleotide exchange factor CaCdc25 (REM and CAT domains) | |
|---|---|---|
| | Dataset A | Dataset B |
| **Data Collection** | | |
| X-ray beamline | BL13-XALOC (ALBA) | |
| Space group | $P2_12_12_1$ | |
| Cell dimensions | $a$ = 49.1 Å<br>$b$ = 107.8 Å<br>$c$ = 197.7 Å | $a$ = 48.6 Å<br>$b$ = 107.2 Å<br>$c$ = 196.9 Å |
| Wavelength (Å) | 0.97915 | 0.97926 |
| Resolution range (Å) | 98.84-3.00 (3.08-3.00) | 43.22-2.45 (2.50-2.45) |
| Total / Unique reflections | 181,241 / 21,838 (19,494 / 2,154) | 190,781 / 38,604 (9,513 / 1,803) |
| Average multiplicity | 8.3 (9.1) | 4.9 (5.0) |
| Completeness (%) | 99.8 (100.0) | 99.9 (100.0) |
| R meas[b] (%) | 51.3 (305.7) | 16.5 (168.5) |
| R pim[c] (%) | 18.0 (98.2) | 7.3 (75.3) |
| $CC_{1/2}$ (%) | 96.9 (41.0) | 99.5 (45.4) |
| Mean $I/\sigma I$ | 7.0 (1.3) | 8.5 (1.2) |
| **Refinement** | | |
| Resolution range (Å) | 47.68 – 3.0 | 43.22 – 2.45 |
| Unique reflections, work/free | 21,833 / 1,068 | 38,543 / 1,953 |
| R work (%) | 30.9 | 19.4 |
| R free[d] (%) | 34.0 | 24.0 |
| Number of: | | |
| Amino acid residues | 799 | 840 |
| Water molecules | – | 144 |
| Average B value (Å$^2$) | | |
| Wilson plot | 72.2 | 45.2 |
| Protein | 68.8 | 54.6 |
| Solvent | – | 52.4 |
| rmsd bond lengths (Å) | 0.003 | 0.002 |
| rmsd bond angles (°) | 0.67 | 0.45 |
| Ramachandran | | |
| Favored (%) | 82.8 | 98.2 |
| Allowed (%) | 10.6 | 1.8 |
| Outliers (%) | 6.6 | 0 |
| Rotamer outliers (%) | 15.5 | 0 |
| Clashscore | 8.8 | 2.2 |
| PDB entry | – | 7NZZ |
| SBGrid Data Bank entry | doi:10.15785/SBGRID/860<br>doi:10.15785/SBGRID/861 | doi:10.15785/SBGRID/859 |

[a]Values in parenthesis correspond to the outermost resolution shell. [b]R meas is the multiplicity independent R factor. [c]R pim is the precision-indicating merging R factor. [d]Calculated using 5% of reflections that were not included in the refinement.

**Table S3. Comparison of the specific activities between different GEFs with Cdc25 homology domains.**

| GEF | Specific activity $(k)^a$ / $10^5$ $M^{-1}$ $s^{-1}$ | ratio $k_{CaCdc25}/k_{GEF}$ |
|---|---|---|
| CaCdc25 | 4[b] | 1 |
| Sos1 | 0.002[c] | 2000 |
| RasGRP1 | 0.02[c] | 200 |
| RasGRP2 | 0.05[c] | 80 |
| RasGRP3 | 0.01[c] | 400 |
| PDZ-GEF1 | 0.8[c] | 5 |
| PDZ-GEF2 | 0.8[c] | 5 |
| Rlf | 0.04[d] | 100 |
| C3G | 0.03[e] | 133 |

[a]As $k_{obs}$ / [GEF]. [b]This work. [c]Specific activity values derived from (Popovic et al., 2013) (only values for which GEFs showed to be efficient were taken account). [d]Value taken from (Popovic et al., 2016). [e]Value taken from (Carabias et al., 2020).

**Table S4. Oligonucleotides used to amplify CaRas1 and the GEF catalytic region of CaCdc25 ORFs.**

| Primer name | Sequence (5' - 3') |
|---|---|
| Ras1-166 Forward | CCGGCCATGGCGCTGCGTGAATACAAAC<br>NcoI A L R E Y K |
| Ras1-166 Reverse | CCGGGGTACC*TTA*GTTGATGTTGCGCACCAG<br>Acc65I N I N R V L |
| Ras1-Full Reverse | CCGGGGTACC*TTA*TACAATGAC<br>Acc65I V I V |
| GEF Forward | CCGGCCATGGGCAATAATACGAGTT<br>NcoI G N N T S |
| GEF Reverse | CCGGGGTACC*TTA*TTTCAGCGAGAAC<br>Acc65I K L S F |

**Table S5. Oligonucleotides used for site-directed mutagenesis of CaRas1 and of GEF catalytic region of CaCdc25.**

| Primer name | Sequence (5' - 3') |
|---|---|
| Ras1-213 Forward | CCAAATCAATAAC***T***A***G***AACAACACTTCTGCAGTCAATGGC<br>Stop |
| Ras1-213 Reverse | GCCATTGACTGCAGAAGTGTTGTT***C***T***A***GTTATTGATTTGG<br>Stop |
| Ras1-C286A-C287A Forward | CGAGTAGCAAATCCAAGAACGGC**GC**T**GC**CGTCATTGTA*TAAG*<br>A A |
| Ras1-C286A-C287A Reverse | C*TTA*TACAATGACG**GC**A**GC**GCCGTTCTTGGATTTGCTACTCG<br>A A |
| GEF-H1234E Forward | GACCTTCGTCTAC**G**A**A**GGCAACCCGGACTATC<br>E |
| GEF-H1234E Reverse | GATAGTCCGGGTTGCC**T**T**C**GTAGACGAAGGTC<br>E |
| GEF-H1234D Forward | GACCTTCGTCTAC**G**ACGGCAACCCGGACTATC<br>D |
| GEF-H1234D Reverse | GATAGTCCGGGTTGCCGT**C**GTAGACGAAGGTC<br>D |
| GEF-H1234A Forward | GACCTTCGTCTAC**GC**CGGCAACCCGGACTATC<br>A |
| GEF-H1234A Reverse | GATAGTCCGGGTTGCCG**GC**GTAGACGAAGGTC<br>A |
